# Sialidases derived from *Gardnerella vaginalis* and *Prevotella timonensis* remodel the sperm glycocalyx and impair sperm function

**DOI:** 10.1101/2025.02.01.636076

**Authors:** Sarah Dohadwala, Purna Shah, Maura K. Farrell, Joseph A. Politch, Jai Marathe, Catherine E. Costello, Deborah J. Anderson

## Abstract

Bacterial vaginosis (BV), a dysbiosis of the vaginal microbiome, affects approximately 30 percent of women worldwide (up to 50% in some regions) and is associated with several adverse health outcomes including preterm birth and increased incidence of sexually transmitted infections (STIs). BV-associated bacteria such as *Gardnerella vaginalis* and *Prevotella timonensis* damage the vaginal mucosa through the activity of sialidase enzymes that remodel the epithelial glycocalyx and degrade mucin glycoproteins. This damage may contribute to adverse health outcomes. However, whether BV-associated glycolytic enzymes also damage sperm has not yet been determined. Here, we show that sialidase-mediated glycocalyx remodeling of human sperm increases sperm susceptibility to damage and adversely affects their function in vitro. Specifically, we report that sperm motility was not adversely affected by sialidase treatment, but desialylated human sperm demonstrate increased susceptibility to agglutination and complement-mediated cytotoxicity as well as impaired transit through cervical mucus. Our results demonstrate mechanisms by which BV-associated sialidases affect sperm survival and function and potentially contribute to adverse reproductive outcomes such as infertility.

## Introduction

Bacterial vaginosis (BV) is a condition characterized by vaginal dysbiosis that affects approximately one third of women globally (Peebles et al. 2019). In some regions in the global South, up to 50% of women have BV (Kenyon et al. 2013). BV is an extremely variable condition, characterized by high diversity in the vaginal microbiome, and clinical symptoms such as irritation, discharge, itching, and fishy odor (Abou Chacra et al. 2022). To more precisely define the composition of vaginal microbiota found in humans, the field has adopted community state types (CSTs) numbered I to V to describe various proportions of different bacterial taxa (Ravel et al. 2011). A healthy vaginal microbiome, or CST I, consists primarily of *lactobacillus crispatus,* which maintains low vaginal pH and inhibits inflammation. A microbiome dominated by *lactobacillus iners*, termed CST III is thought to be an intermediary microbiome between a healthy and disease state. CST IV (i.e. BV) is characterized by a relative lack of lactobacilli and the presence of dysbiotic anaerobic vaginal bacterial species such as *Gardnerella vaginalis* and *Prevotella timonensis*. BV is associated with spontaneous preterm birth, increased risk of preclinical pregnancy loss, and the acquisition of sexually transmitted infections (STIs) (Howe et al. 1999; Ralph et al. 1999; Cauci and Culhane 2011; van Oostrum et al. 2013; Armstrong and Kaul 2021; Abou Chacra et al. 2023; Mohanty et al. 2023, Hadhoum et al 2025.). In addition, emerging evidence suggests a link to infertility and pre-eclampsia, as well as adverse neonatal outcomes even for full term infants (Denney and Culhane 2009; van Oostrum et al. 2013; Dingens et al. 2016; Kindschuh et al. 2024; Ravel et al. 2021).

To date, the mechanisms by which BV increases the risk of adverse reproductive outcomes are poorly understood (Ugwumadu 2002; Mania-Pramanik et al. 2009; Elovitz et al. 2019; Gudnadottir et al. 2022); several have been proposed, including higher vaginal pH, an inflammatory vaginal milieu, epithelial damage and altered innate immunity (Cauci et al. 2008). Recent findings have implicated sialidases, enzymes that cleave sialic acid residues from glycoproteins and glycolipids, in BV-associated adverse reproductive outcomes. Several BV-associated bacteria such as *Gardnerella vaginalis and Prevotella timonensis* can produce high levels of sialidases, and sialylation of the vaginal epithelium and mucus is markedly lower in women with BV than those with a healthy microbiome (Agarwal et al. 2023; Segui-Perez et al. 2024). In particular, elevated sialidase activity at 12-weeks gestation has been correlated with a higher risk of early spontaneous preterm birth and miscarriage, with a stronger correlation than for BV generally (Cauci et al 2008). BV-associated sialidases have been shown to degrade the vaginal glycocalyx and mucus (Pelayo et al 2024; Agarwal et al 2023). While BV-associated degradation of the vaginal mucosa certainly contributes to clinical symptoms of BV (Roselletti et al. 2020; Cheu et al. 2022), the mechanistic connection between BV-associated vaginal changes and adverse reproductive outcomes, which primarily occur in the upper female reproductive tract (FRT), remain unclear. Sperm migrate from the vagina to the upper FRT and it has been postulated that abnormal sperm play a role in female reproductive disorders (Mahdavinezhad et al. 2022). In addition, recent evidence suggests that sperm can modulate the uterine immune milieu which may have an impact on the timing of gestation (Green et al. 2020; Katila et al. 2001; Schjenken et al 2021). Accordingly, the purpose of this study was to determine whether BV-associated sialidases derived from *Gardnerella vaginalis and Prevotella timonensis* affect the human sperm glycocalyx and impact sperm function. The sialylated sperm glycocalyx protects them from the female immune system, facilitates their passage through cervical mucus, and prevents premature capacitation, a process necessary for fertilization (Tecle and Gagneux 2015; Fliniaux et al. 2022). It is estimated that mature sperm carry approximately 30 million sialic acid molecules per cell (Ghaderi et al. 2011, PNAS). Thus, BV-associated sialidases may adversely affect sperm migration, immune signaling, and fertilization capacity.

We used three recombinant sialidases, GvNanH2, PtNanH2 and PtNanH1, derived from clinically relevant strains of *Gardnerella vaginalis* (*Gv*) and *Prevotella timonensis (Pt)*, to study the effects of BV-associated sialidases on human sperm. These enzymes reportedly have different activities: the NanH2 enzymes cleave both *N-* and *O-*linked glycans and play an important role in degrading the protective sialoglycan mucus barrier lining the lower female reproductive tract, whereas the NanH1 enzyme demonstrates reduced enzymatic activity including towards sialic acids linked to human MUC5B and other mucin substrates (Agarwal et al. 2023, Pelayo et al 2024). We employed lectin flow cytometry and zeta potential to characterize the changes to surface glycan moieties and charge on the remodeled sperm glycocalyx after sialidase treatment. We next investigated the effect of desialylation on sperm motility, the ability of sperm to traverse through cervical mucus, and on spontaneous and antibody/lectin-induced sperm agglutination. Lastly, we investigated the susceptibility of desialylated sperm to complement-mediated cytotoxicity. Our data provide strong evidence that BV-associated sialidases adversely affect several sperm functions, potentially contributing to the adverse reproductive outcomes such as subfertility and preterm birth that are associated with bacterial vaginosis.

## Results

### Sialic acids are depleted from the sperm glycocalyx after exposure to recombinant BV sialidase, exposing cryptic epitopes and reducing negative surface charge

Sperm are rich in sialylated cell surface proteins that may be susceptible to hydrolysis by BV-associated sialidases. Accordingly, to investigate BV-associated damage to sperm, we produced recombinant, low endotoxin GvNanH2 as described previously (Robinson et al. 2019, **Fig S1 – S2**). GvNanH2 is a truncated variant of a sialidase derived from *Gardnerella vaginalis.* Of note, the *Prevotella* species do produce endotoxin in the vaginal tract; in our study, *Prevotella* enzymes were produced in BL21 E.coli, and as a result likely contain endotoxin (Aroutcheva et al. 2009). Controls to determine the effects of endotoxins were incorporated into the experiments as described below. To ensure that our experiments employed physiologically relevant amounts of sialidase, we used a range of activity levels of enzyme derived from cervicovaginal swabs obtained from women with BV (0 to 2 uM 4MU/min sialidase activity, mean = 0.5) from Agarwal et al. 2023, Figure 1B. Given those 4-MU activity levels from vaginal swabs, we used 0.88 and 0.11 uM 4-MU/min/uL of enzyme activity to represent BV-like conditions and standardize our assays. To confirm that BV-associated sialidases desialylated sperm-surface glycans, sperm were collected from healthy, reproductive aged men, isolated by density gradient, and treated with either recombinant GvNanH2, AUS (a commercially available sialidase derived from *A. ureafaciens*, positive control), or no treatment (negative control). Sperm were incubated in non-capacitating conditions. Then, we assessed removal of terminal sialic acids with lectin flow cytometry using SNA-Cy5 and MAL II-biotin to measure a decrease in 2,6 and 2,3-linked terminal sialic acids, respectively (**Fig 1A**). The untreated controls demonstrated the presence of 2,3 and 2, 6 linked sialic acids on the sperm surface (**Fig 1B**, **1C).** This corresponds to results from previous studies using lectin microarrays to profile the sperm surface glycome (Xin et al. 2014). For SNA-Cy5 staining, the direct conjugate enabled a direct comparison of the activity of AUS and GvNanH2. AUS reduced the signal by roughly 50% relative to the control, while GvNanH2 reduced it by approximately 15% (**Fig 1C**). Similarly, a significant reduction was observed in the staining for MAL-II, though due to a secondary amplification step, a linear relationship to compare AUS and GvNanH2 was not possible. Although not statistically significant, a three-fold decrease in signal was observed with NanH2 compared to AUS (**Fig 1B**).

**Figure 1:**
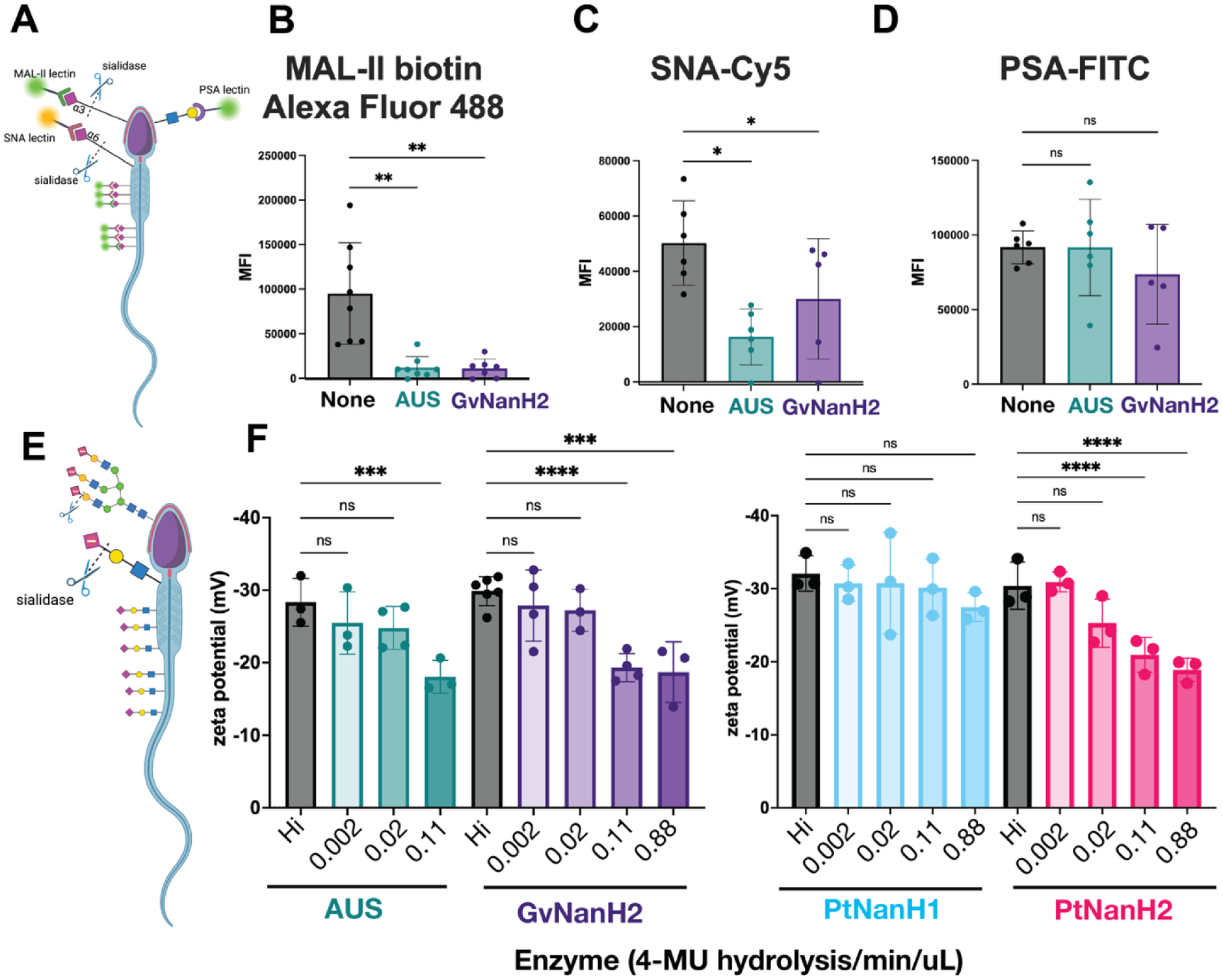
BV-associated sialidases remove surface sialic acid from sperm. A) Schematic describing lectin flow cytometry and representative glycan epitopes. B, C, D) Sperm were exposed to sialidases (AUS 0.11 4-MU hydrolysis/min/uL or GvNanH2 0.88 4-MU hydrolysis/min/uL) for 1 hour, fixed in formalin, and stained with MAL-II-biotin/Neutravidin-Alexa Fluor 488, SNA-Cy5, or PSA-FITC respectively. Each point represents the median fluorescence intensity (MFI) from one donor. Statistics represent a mixed model comparison with post-hoc Holm-Šidák tests. E) Schematic representing change in zeta potential after removal of negatively charged sialic acid on sperm surface. F) Sperm were treated with sialidase at various doses for 1 hour, and zeta potential was measured via dynamic light scattering. Each measurement represents the mean of seven technical replicates, and statistics represent a mixed model comparison with post hoc Holm-Šidák tests. * indicates p ≤ 0.05, ** p ≤ 0.01, ***p ≤ 0.001, and **** p ≤ 0.0001. Hi indicates heat-inactivated at the highest dose of enzyme (0.88 for PtNanH1, PtNanH2, and GvNanH2, and 0.11 for AUS).

Sperm, being foreign to the female reproductive tract, have immunogenic epitopes capped with terminal sialic acids that function in processes such as the acrosome reaction and sperm-egg interaction (Ma et al 2016). To determine if the glycan moieties underlying terminal sialic acids were exposed or degraded after sialidase treatment, we used lectin flow cytometry with pea savitum agglutinin (PSA), a lectin which binds *N-*acetyllactosamine residues and is a marker for the acrosome reaction (**Fig 1A**). In untreated (control) sperm, robust PSA staining was observed (**Fig 1D**). In both sialidase treatment conditions (AUS and NanH2), sperm PSA staining was unchanged from that of the control (**Fig 1D**). This indicates that sialidase treatment does not result in a premature acrosome reaction under these conditions.

Sperm surface sialic acids contribute to a negative surface charge. To determine if the surface charge of sperm is modified in response to sialidase treatment, the zeta potential of sperm cultures was measured after treatment with different doses of enzyme. First, we optimized the diluent for the assay and confirmed the effect of different culture conditions on the measurement (**Fig S4A**). A decrease in sperm surface charge was observed when whole semen was treated, though the magnitude of the change observed was lower than when washed sperm were treated (**Fig S4B**). It is possible that sialylated glycoproteins in seminal plasma protect the sperm surface from the full force of the sialidase enzymes through a competition mechanism. Next, we measured the change in zeta potential to compare the same doses of recombinant NanH2 and AUS and found that effects were similar (**Fig S4C**). Finally, we used a series of doses of GvNanH2, PtNanH2, PtNanH1, and AUS sialidase and measured the zeta potential of the sperm. We observed a dose dependent decrease in the magnitude of the negative surface charge after a 1-hour incubation for AUS, PtNanH2, and GvNanH2, though we did not observe a significant decrease with PtNanH1 (**Fig 1F**). This indicates that glycans that cannot be cleaved by PtNanH1 are a major contributor to the negative surface charge of sperm. Based on the limited data on the substrate specificity of PtNanH1 and PtNanH2, we surmise that these sialylated proteins may be similar to mucin glycan substrates such as MUC5B in structure or accessibility. We also measured the donor-to-donor variance in the biological replicates and found that there was some variance in how the sperm from different donors were affected by the sialidases, though overall trends in the data were retained (**Fig S4D**).

### Desialylated sperm are more susceptible to agglutination

Agglutination is a phenomenon driven by cell motility and surface charge wherein motile sperm collide and stick to one another, forming large, aggregated networks. We investigated the propensity of highly motile sperm to agglutinate after sialidase treatment. We hypothesized that by reducing the negative charge, sperm desialylation would make the sperm stickier and prone to agglutination.

To measure this, washed sperm were treated with different doses of PtNanH2 sialidase and observed over time. We observed variable degrees of spontaneous sperm agglutination under these conditions, largely because agglutination is a concentration dependent phenomenon that relies on cell-cell collisions. The variable characteristics in these spontaneous agglutinates included size and shape of agglutinates (**Fig S9, S10**). Next, we tested whether sperm desialylation affects antibody- or lectin-induced sperm agglutination. To quantify this effect, we performed a kinetic agglutination assay on sperm treated with sialidase or heat inactivated enzyme (negative control), followed by Galectin-1, the Human Contraception Antibody (HCA), or Pea savitum agglutinum (PSA) **(Fig 3A).** These entities were selected due to their importance in reproductive biology. Galectin-1, an endogenous lectin, is involved in binding STI pathogens and is expressed in FRT tissues. HCA, a sperm-binding monoclonal antibody derived from the blood of an infertility patient, is part of a contraceptive product that is poised to enter Phase II clinical trials (Anderson et al. 2020). Finally, PSA is a marker for the acrosome reaction, a process crucial to sperm-egg interaction. These three entities bind distinct glycans--HCA binds a male reproductive tract specific, but largely uncharacterized sperm *N*-glycoprotein; Galectin-1 binds repeat *N*-acetyllactosamine units, and PSA binds terminal mannose residues. None of these three entities bind sialic acid moieties specifically, though they may all be tolerant of sialylated glycans.

We optimized the concentrations of the agglutinating antibodies and lectins, selecting low concentrations that were weakly agglutinating to distinguish the effect of sialidases treatment. We found that sperm agglutinated significantly faster in response to each of the three agglutinating entities after treatment with either AUS or GvNanH2 sialidase (**Fig 3B**, **3E**). We also tested the *Prevotella* sialidases using HCA to assess whether the specificity differences between PtNanH1 and PtNanH2 would have different functional implications related to cell adhesion and agglutination. We found that PtNanH1 did not increase agglutination speed, while PtNanH2 did, similar to GvNanH2 in a dose dependent manner (**Fig 3C**, **3F**). This indicates that sialylated glycans that are cleaved by PtNanH2 but not by PtNanH1 have important roles in preventing agglutination and cell-cell sticking. Given reports in the literature that demonstrate an impact of endotoxin (lipopolysaccharides, LPS) on sperm capacitation, we used polymyxin-B, an antibiotic that neutralizes the effects of LPS, to isolate the role of sialidases and exclude contributions from endotoxin in our assays (Bhagwat et al. 2025). Sperm agglutination assays were repeated in the presence of sialidase pre-treated with polymyxin-B, and there was no difference in activity, indicating that the effect on sperm agglutination can indeed be attributed to sialidase activity (**Fig 3D**). In addition, for PSA, flow cytometry data indicated that surface staining is unchanged in response to sialidase treatment, suggesting charge-mediated repulsion in preventing agglutination, rather than enhanced binding of underlying glycan epitopes in treated sperm (**Fig 1D).** In sum, removal of surface sialic acids from sperm increased their propensity to agglutinate spontaneously and in the presence of several distinct lectins and antibodies.

### Desialylated sperm are more susceptible to complement-mediated immobilization and cytolysis

Sialic acids prevent aberrant complement activation by binding several complement inhibitors including Factor H (Blaum et al. 2015). Both complement and complement inhibitors can be found in human cervical mucus and seminal plasma (Price and Boettcher 1979; Petersen et al. 1980) suggesting that complement biology may be important to fertility and reproduction, though specific mechanisms are not yet fully understood. For instance, complement proteins may serve as a bridge between sperm and egg (Anderson et al. 1993), and some antisperm antibodies demonstrate potent complement-dependent immobilization and cytolysis of sperm (Baldeon-Vaca et al. 2021).

We used the complement-mediated sperm immobilization assay to determine the potential role of sialidase enzymes in spontaneous- and lectin-mediated complement activation. We first investigated whether sialidase treatment alone affected sperm motility. To isolate the measurement of sperm motility from the spontaneous agglutination we sometimes observed, these motility assays were done at low concentrations of sperm at which agglutination was not observed. We did not observe a significant effect of any of the tested sialidases on sperm motility through 60 minutes, after which time we observed a slight decrease in sperm motility in some donors (**Fig 2A**, **2C).** In addition, we plotted average sperm motility parameters that relate to capacitation, including curvilinear velocity, amplitude of lateral head displacement, and linearity over time to determine if sialidase treatment impacted these parameters rather than total motility. We found that, under non-capacitating experimental conditions for 90 minutes, sialidases do not impact hyperactivation motility parameters associated with sperm capacitation (**Fig 2B-D, 2G-H**). Next, we examined the effect of sialidases on sperm motility in the presence of complement. Addition of GvNanH2 and PtNanH2 with complement significantly reduced sperm motility compared to all three negative controls (no enzyme treatment, heat inactivated complement, addition of compstatin, a C3 complement inhibitor) (**Fig 4A**, **4B**). We found that sperm exposure to PtNanH1 did not mediate complement immobilization, which indicates that the sialylated glycans that are cleaved by the NanH2s, but not cleaved by PtNanH1 play another critical role on the surface of sperm in protection from complement (**Fig 4B**, **4C**). Finally we confirmed that complement-mediated sperm immobilization is a surrogate for cell lysis with trypan blue staining (**Fig S10**). After complement treatment of sialidase-treated sperm the majority of immotile sperm stained positive for trypan blue, indicating that they were lysed (**Fig S10**). The majority of lysed sperm were present in agglutinates, which could indicate that sperm become sticky after lysis, or that agglutination facilitates complement activation on the surface of sperm due to increased surface area of the agglutinate (Dalia et al 2011). We confirmed that complement activation could be abrogated using sialidase inhibitors, including DANA, a competitive inhibitor, and Zanamivir, a sialidase inhibitor used for treatment of flu. PtNanH2 was susceptible to inhibition by Zanamivir, while GvNanH2 was susceptible to DANA (**Fig 4D**). In addition, we explored antibody-mediated complement lysis, and found that sperm were more prone to complement lysis through the classical complement pathway in the presence of sialidase (**Fig S5**). In sum, these data provide evidence that surface sialic acid moieties protect sperm from complement-mediated lysis. Finally, given prior reports that endotoxin (which can be found in recombinant bacterial enzymes) may itself impact sperm health, we repeated complement assays in the presence of 100 ug/mL polymyxin B to neutralize endotoxin (Bhagwat et al. 2025). We observed a small (10.5%) but statistically significant motility difference between the polymyxin pre-treated GvNanH2 and GvNanH2 alone (**Fig 4E**). Thus, endotoxin may account for some of the observed sperm immobilization in our assays. However, in the presence of polymyxin B, sialidase-treatment resulted in an average motility reduction of approximately 40% (GvNanH2) and 70% (PtNanH2), compared to polymyxin only controls (**Fig 4E**). The differences in motility between the sialidase-treated and no treatment group in the presence of polymyxin remain highly significant indicating that sialidase treatment does account for the increased sensitivity to complement-mediated immobilization.

**Figure 2:**
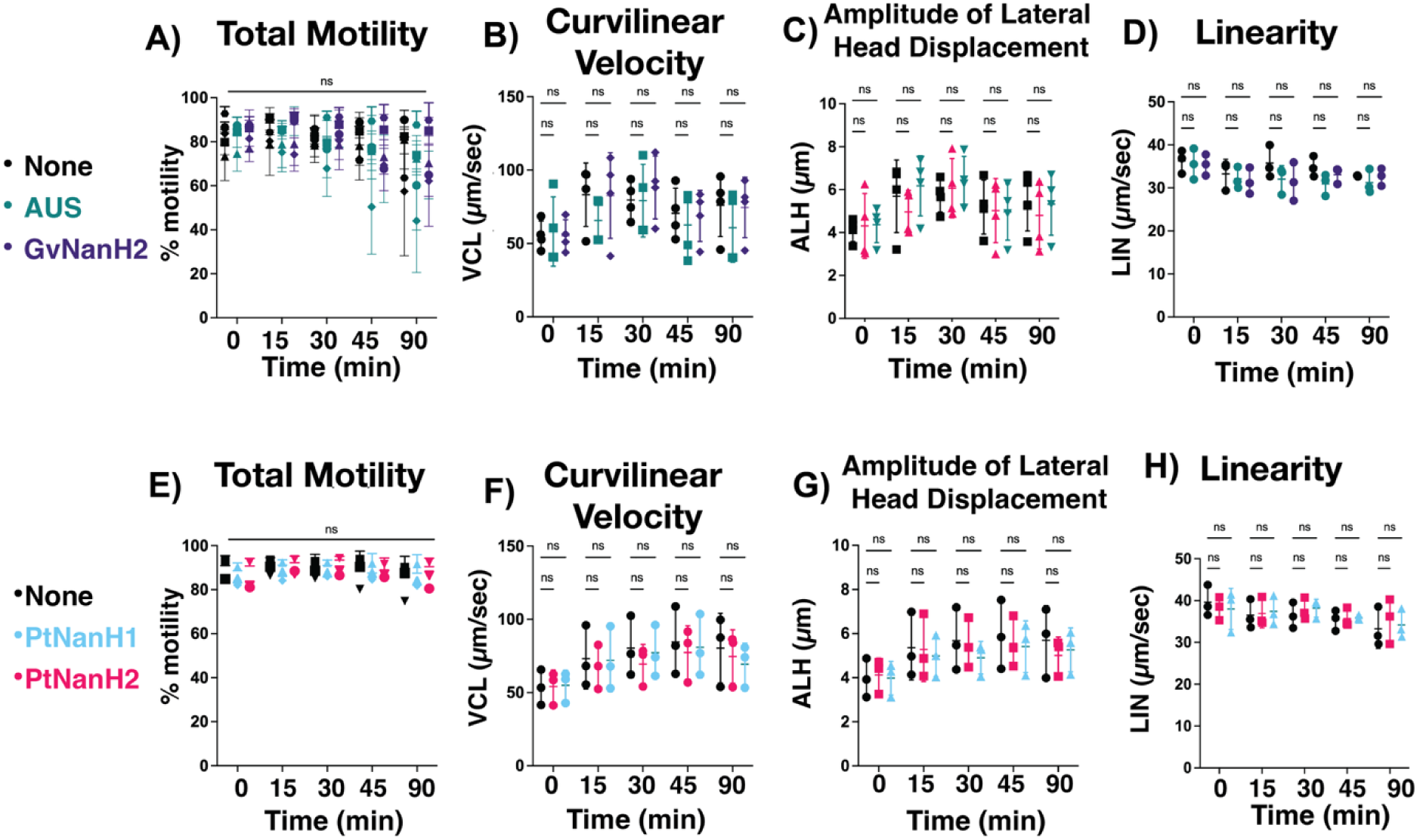
Effect of sialidase treatment on sperm motility. A) Motile sperm were isolated, and motility was measured via computer assisted sperm analysis (CASA) after sialidase treatment (0.88 4-MU hydrolysis/min/uL GvNanH2 and 0.11 4-MU hydrolysis/min/uL AUS). Statistics represent a repeated measures comparison, and neither time nor enzyme condition were statistically significant. E) same as A, except using PtNanH1 and PtNanH2 at 0.88 4-MU hydrolysis/min/uL. B, C, D) Average curvilinear velocity (VCL), Amplitude of lateral head displacement (ALH), and Linearity (LIN), respectively, as measured by CASA on the motile sperm population after treatment with enzyme (0.88 4-MU hydrolysis/min/uL GvNanH2 and 0.11 4-MU hydrolysis/min/uL AUS). F, G, H) Same as B, C, D, except using PtNanH1 and PtNanH2 at 0.88 4-MU hydrolysis/min/uL. Each measurement represents the mean of four technical replicates. Statistics represent mixed model comparisons with post-hoc holms Sidak tests, and neither time nor enzyme condition were a significant source of variation for LIN values. For ALH and VCL values, only time was statistically significant (p<0.01).

**Figure 3:**
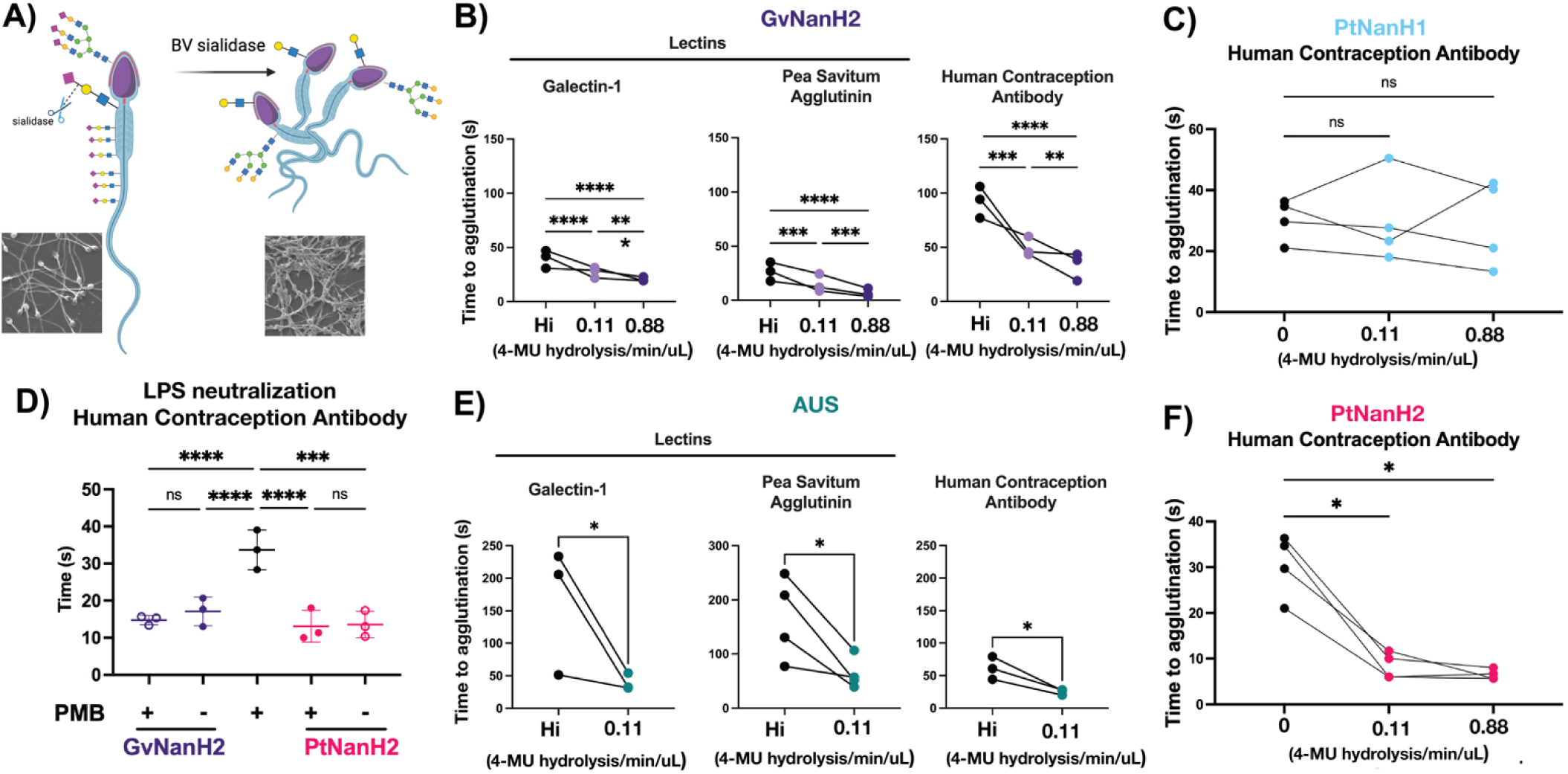
Desialylation results in faster sperm agglutination. A) Schematic and representative images representing sperm agglutination after sialidase treatment. B C, E, F) Time to agglutination in the presence of different lectins and glycan-binding antibodies after treatment with GvNanH2, PtNanH2, AUS and PtNanH1, respectively. Galectin-1 was used at a concentration of 10 µg/mL, PSA was used at a concentration of 0.05 µg/mL, and HCA was used at 5 µg/mL. D) Time to agglutination in the presence of 0.88 4-MU hydrolysis/min/uL of enzyme in the presence and absence of polymyxin B (PMB). Each measurement represents the mean of three technical replicates. Statistics represent repeated measures ANOVA with post hoc Holm-Šidák tests for GvNanH2, PtNanH2, and PtNanH1 (1B, 1C, 1D,1F), and unpaired t-tests for AUS (1E). * indicates p ≤ 0.05, ** p ≤ 0.01, ***p ≤ 0.001, and **** p ≤ 0.0001. Hi indicates heat-inactivated enzyme at the highest concentration.

**Figure 4:**
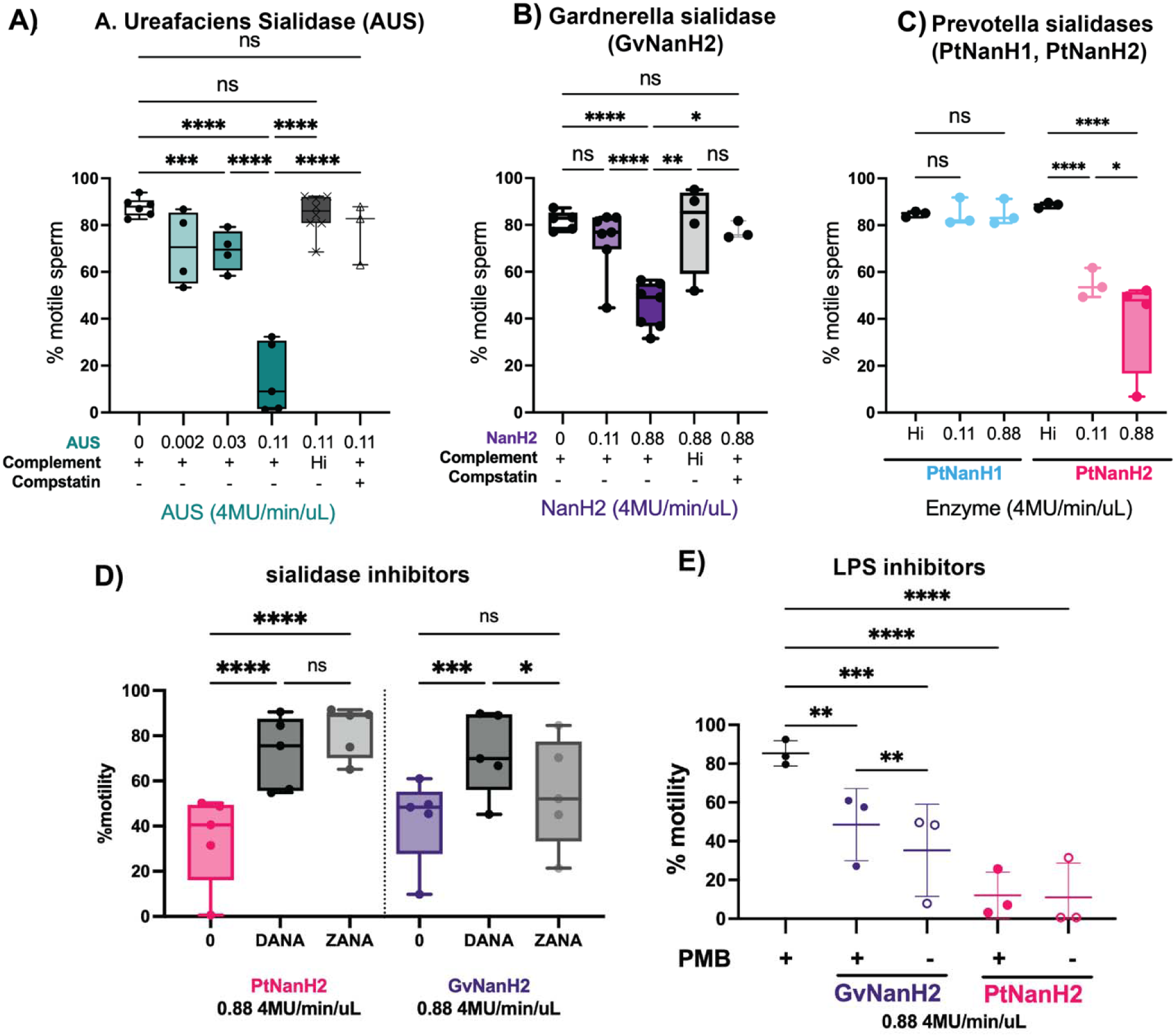
Effect of sialidase treatment on susceptibility to complement-mediated immobilization. A, B, C) Sperm motility was measured after treatment with sialidase in the presence of complement. Heat-inactivated complement, or 125 uM compstatin, a human C3 inhibitor, were used as negative controls. Statistics represent mixed model comparisons with post-hoc Holms-Sidak tests. Hi indicates heat inactivated. D) The effect of sialidase inhibitors on complement immobilization of sperm with GvNanH2 and PtNanH2. DANA indicates N-acetyl-2,3-dehydro-2-Deoxyneuraminic Acid, and ZANA indicates Zanamivir. E) The effect of LPS inhibitor polymyxin B (PMB) on complement immobilization of sperm with GvNanH2 and PtNanH2 at 0.88 4-MU hydrolysis/min/uL of activity. Each measurement represents the mean of three technical replicates. Statistics represent a mixed model with post-hoc Holms-Sidak tests. * indicates p ≤ 0.05, ** p ≤ 0.01, ***p ≤ 0.001, and **** p ≤ 0.0001.

### Desialylated sperm are less able to transit through human cervical mucus

Human cervical mucus varies in physical properties and composition throughout the menstrual cycle and is rich in negatively charged mucins. Sperm must transit across this mucosal barrier to access the upper reproductive tract and fertilize an egg. To determine how sperm transit through mucus may be affected by sialidase treatment, we assessed numbers of sperm observed at progressive distances through a capillary tube filled with ovulatory human cervical mucus. We started with a model that simulates sperm transit through the vagina and cervix after intercourse (**Fig** 5A). Motile sperm were isolated, resuspended at a concentration of 80 million/mL in 50% seminal plasma, and placed into a capillary tube to measure migration through a small region (0.5 cm) of GvNanH2 sialidase or media (control), followed by midcycle cervical mucus (**Fig** 5A). We found a significant reduction in the numbers of total sperm that were able to transit through the mucus after exposure to sialidase (**Fig** 5B).

**Figure 5:**
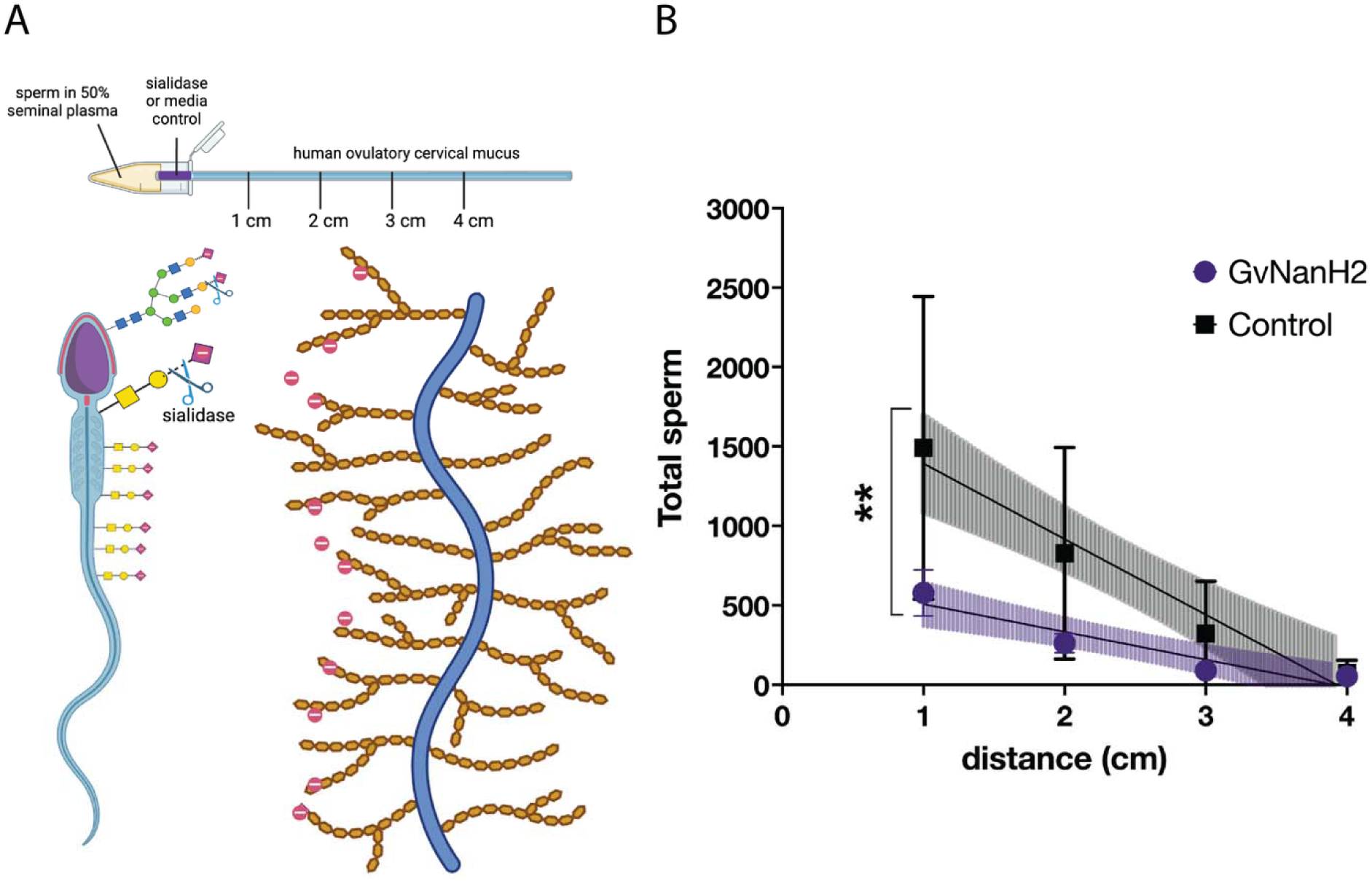
Sperm transit through cervical mucus is reduced after desialylation. A) Schematic depicting experimental setup and mucus-sperm interaction model. Sperm were incubated with seminal plasma, then inserted into a capillary tube containing sialidase at the entrance and ovulatory cervical mucin. B) Total sperm were quantified at each distance in the capillary tube containing ovulatory cervical mucus, and either 0.5 4-MU hydrolysis/min/uL GvNanH2 sialidase or media at the entrance. Statistics represent an unpaired test comparison of the slopes of the lines derived from the four points. Data represent three biological replicates with either two or three technical replicates depending on donor volume.

Next, we measured different methods of sialidase exposure since the localization of the bacterial enzyme in the cervix is unknown. We mixed human midcycle cervical mucus with recombinant sialidase (amount equivalent to 0.5 μM/min 4-MU hydrolysis activity) or media (control) and counted the number of sperm that penetrated to various depths of the capillary tube after 90 minutes. Pre-treatment of mucus with sialidase initially facilitated sperm transit, but the sperm became immotile as they traversed the sialidase-containing mucus (**Fig S7**). While the numbers of sperm found in mucus increased in the sialidase-containing mucus, the percent motile sperm decreased, which was the opposite of the trend observed in untreated mucus (**Fig S7**). This may be due to more efficient complement lysis of the sperm exposed to sialidase during transit through the cervical mucus. Next, we tested treatment of sperm with sialidase prior to mucus exposure and found that this also reduced their transit through cervical mucus (**Fig S7**). We also tested these phenomena with and without seminal plasma, which contains several proteins known to coat the sperm surface (**Fig S6**). We found that seminal plasma protects sperm from the immobilization observed in sialidase-containing mucin, perhaps due to abundant seminal sialoglycoproteins in seminal plasma (**Fig S6**). Finally, we also compared sperm transit through human cervical mucus to a commonly used commercially available mucin model, bovine submaxillary mucin. In the absence of seminal plasma, a similar pattern was observed; however, in the presence of seminal plasma, human cervical mucus was much more permissive to sperm transit (**Fig S8**). This indicates some specific interaction between components of human cervical mucin and seminal plasma that is not recapitulated with bovine mucin. In sum, our data indicate that BV-associated sialidases affect sperm transit through human midcycle cervical mucus, both by altering the sperm’s forward progression in mucus, and by altering the properties of the mucus itself.

## Discussion

Our study demonstrates that sialidases derived from BV-associated bacteria alter the sperm glycocalyx by removing terminal sialic acid residues. Furthermore, it provides evidence that the unperturbed sperm glycocalyx facilitates the transit of sperm through cervical mucus and likely protects sperm from immune effectors such as complement in the female reproductive tract. Desialylated sperm were highly susceptible to complement-mediated damage and agglutination, and demonstrated impaired penetration through midcycle cervical mucus. Prior to our work, most studies on the effects of BV-associated sialidases were limited to the vaginal mucosa (Agarwal et al. 2023; Segui-Perez et al. 2024), and the importance of sperm surface sialic acids in bacterial infection and capacitation (Khosravi et al. 2016). Studies related to the effects of sialidases on sperm were mostly conducted in animal models which supported the idea that exposure to sialidases affects sperm function in the reproductive tract (Ma et al. 2016; Fernandez-Fuertes et al. 2018).

Several orthogonal lines of evidence support the idea that BV-associated sialidases may damage sperm and affect fertility: 1) *G.vaginalis* is found at a higher frequency in semen from infertile men (Andrade-Rocha 2009) and oligozoospermic seminal plasma contains higher levels of free sialic acid relative to normozoospermic samples (Menevse et al. 2022), 2) other bacterial conditions that involve glycolytic enzymes are mechanistically linked to male infertility (Khosravi et al. 2016), 3) multiple studies with sperm from various other mammalian species have pointed to the importance of the glycocalyx in protecting from innate immunity in the FRT (Fliniaux et al. 2022), 4) the semen microbiome is associated with sperm health (He et al. 2024; Osadchiy et al. 2024) and 5) natural reproductive processes, such as capacitation, the acrosome reaction and sperm-egg fusion, are sialidase-dependent and rely on the timely exposure of relevant surface glycans (Feng et al. 2016). Finally, several animal models demonstrate that sialic acid moieties on sperm can impact immune cells and endometrial reproductive processes (Toshimori et al. 1991; Froman and Thursam 1994; Machado et al. 2014; Álvarez-Rodríguez et al. 2020; Batra et al. 2020; Schjenken et al. 2021; Donnellan et al. 2023; Li et al. 2024). In summary, human sperm are an underappreciated target for bacterial sialidase enzymes associated with BV; premature desialylation likely impacts sperm function and the immunology of the female reproductive tract.

We found that desialylated sperm provoke complement activation resulting in sperm immobilization and death. In addition, sperm exposure to endotoxin may synergize with sialidase treatment to further sensitize sperm to complement immobilization (Bhagwat et al. 2025). The premature death of sperm in the vaginal and cervical environment may explain the association of BV with infertility (Ravel et al. 2021). The appropriate activation of the complement cascade is important to several reproductive processes, including fertilization and early pregnancy, and disturbances resulting in aberrant complement activation are associated with adverse pregnancy outcomes including preterm birth (Anderson et al. 1993; Riley-Vargas et al 2005; Lynch et al. 2011; Lynch et al 2008; Regal et al. 2015). In particular, higher levels of complement components C3b and C5, and mediators such as mannose-binding lectin have been found in vaginal swabs from pregnant women who deliver preterm compared to women who delivered term (Chan et al. 2022). Given that sperm transit through the lower reproductive tract to the upper tract, we hypothesize that desialylated sperm may initiate and carry some of these aberrant complement cascades into the upper tract prior to fertilization (Schjenken et al. 2021; Denny et al 2012). Early processes in pregnancy, including implantation and placentation, are inflammatory in nature, but overactivation of these pathways results in pathology; perhaps desialylated sperm are partially responsible for the imbalance. (Tantengco and Menon 2022; Norwitz et al. 2001; Norwitz 2006).

We found that sialidase did not affect sperm motility including hyperactivated motility. Given that this study focuses on the impacts to sperm that may occur in the vaginal tract or the cervix, we did not study impacts on sperm under capacitating conditions, which typically are considered to represent the uterine environment. Given that sperm do alter their surface sialylation during capacitation, further studies to clarify whether endogenous desialylation is required for capacitation or whether bacteria mediated desialylation impacts capacitation could be informative.

Next, we found that desialylated sperm are more prone to agglutination, probably due to a reduction in the surface charge that removes inherent cell-cell repulsion. Our data indicate that the surface charge of sperm is largely due to the presence of specific sialylated glycans (those cleaved by NanH2s but not PtNanH1), which are crucial for cell-cell repulsion. Enhanced agglutination may mean that contraceptive antibodies and lectins will be more efficacious in women with sialidase-positive BV (Tsuji et al. 1988; Baldeon-Vaca et al. 2021). The process of agglutination is entirely dependent on sperm concentration, due to its dependence on cell-cell collisions. Accordingly, the extent and nature of spontaneous agglutinates varies and it remains unclear how best to accurately measure spontaneous agglutination in a context that would reflect in vivo conditions. Given that several leading clinical contraceptive candidate antibodies as well as a few anti-HIV antibodies rely on glycan binding, it will be important to investigate how women with sialidase-positive BV respond to these topical glycan-binding drug products (Anderson et al. 2017). We believe the enhanced agglutination is due to reduced surface charge of the sperm, rendering them “stickier. Several groups have proposed that sperm may carry STI pathogens into the upper tract via cell-cell sticking, and the changes in surface charge due to vaginal sialidases may exacerbate this phenomenon (Young et al. 2019).

We found that desialylated sperm traverse less successfully through ovulatory cervical mucus. This supports data from cervical mucus in the sheep model, where differences in sialic acid content in cervicovaginal tissue and mucus were associated with impaired fertility (Abril-Parreño et al. 2022). The human cervical mucus penetration assay was originally designed as a fertility test, so the impaired ability of desialylated sperm in this assay suggests that sialidases may impact fertility (Hull et al. 1984). Studies on the nature of ovulatory human cervical mucus are hindered by the difficulty of sample collection, particularly from women with untreated bacterial vaginosis and in the absence of hormonal birth control, which can modify mucus characteristics (Moncla et al. 2016). It is known that women with BV produce more mucus, perhaps to counteract the bacterial metabolism of mucin glycoproteins observed in BV (Moncla et al. 2016). However, it is unknown whether sialidases remain in the vagina or ascend into the cervical environment and upper reproductive tract in women with sialidase-positive BV.

Given that mucin-like glycans contribute significantly to the surface charge of sperm and mediate protection from complement lysis and agglutination, it is important to understand which sialylated proteins mediate these functions, and to more granularly define the specificity differences that exist between the NanH2s and PtNanH1. Much work remains in understanding the roles of sialylated glycoforms of proteins on the sperm surface, and even how some sialylated proteins, such as the semenogelins and beta defensin 126, which are abundant mucin-like proteins critical for semen liquefaction and mucus penetration, respectively, adhere to the sperm surface (Luo et al 2023, Diao et al 2014; Tollner at al 2011, Sci Trans, Med, Dorin and Barratt 2014 Hum Reprod.)

This work has some limitations. The fertility status of our sperm donors was unknown; we did not collect samples from infertile or subfertile men, so we cannot make assertions about male infertility outside the FRT, or how bacterial sialidases may exacerbate pre-existing male infertility. In addition, we did not screen sperm donors for BV-associated bacteria, though sperm samples with poor motility were excluded. In addition to sialidases, several other glycolytic enzymes including amylases and fucosidases are produced by BV-associated bacteria that we have not yet explored. Due to limitations in human cervical mucin availability, we did not assay the *Prevotella* sialidases’ impact on sperm movement through mucus. Finally, the diversity in bacterial vaginosis cases should also be further considered, given that molecular profiling has revealed varying presentations of vaginal flora (Savicheva et al. 2023). *Gardnerella vaginalis and Prevotella timonensis*, which produce the sialidases of focal interest in this study, are a primary cause of BV (Morrill et al. 2020). However, it should be noted that not all genotypes of G. *vaginalis* produce sialidase, and this may cause variation in clinical presentation of BV cases. Furthermore, other species such as mycoplasma, express additional sialidases (Hardy et al. 2017). Clearly, in adverse reproductive outcomes associated with BV, there is also a role of the damage to the FRT. This work did not study the interactions between damaged FRT and sperm. The *in vitro* environment may not recapitulate certain aspects of physiology, such as fluid flow, muscle contractions, as well as the menstrual cycle and hormone signaling on sperm. In addition, we did not assess interactions between sperm and uterine immune cells which may be impacted by desialylation, such as targeting of sperm by neutrophils via NETosis, sperm phagocytosis mediated by uterine macrophages, or sperm immunomodulation of uterine NK cells.

In summary, we demonstrate that BV-associated sialidases increase sperm susceptibility to damage in the FRT and impede their penetration through cervical mucus, providing evidence that these mechanisms may play an underappreciated role in adverse reproductive outcomes. Understanding aberrant immune damage to sperm in bacterial vaginosis raises new questions about how sperm may modify the immune milieu of the female reproductive tract in both a eubiotic and dysbiotic condition. These findings highlight the need for further investigation of the role of sperm in decidualization and ascending inflammation in the upper reproductive tract, as well as a better foundational understanding of important processes of early pregnancy in humans.

## Materials and Methods

### Materials

#### Enzymes and substrates

AUS refers to Neuraminidase A (New England Biolabs P0722L). 4-MU Sialic acid refers to 2′-(4-methylumbelliferyl)-α-D-*N*-acetylneuraminic acid sodium salt hydrate (Sigma Aldrich 69587). GvNanH2 refers to truncated recombinant *Gardnerella vaginalis* derived NanH2, as described below. PtNanH1 and PtNanH2 refer to enzymes derived from *Prevotella timonensis*, isolated and produced as described in Pelayo et al 2024.

#### Bacterial strains, plasmids and DNA

ClearColi bacteria strains that produce low endotoxin protein were a gift of the Dempsey lab at BUMC. The GvNanH2 plasmid was a gift of the Lewis lab at UCSD.

#### Production of recombinant NanH2 (GvNanH2)

The GvNanH2 plasmid contains a functional, soluble, His-tagged, truncated form of NanH2, a sialidase expressed by *Gardnerella vaginalis*, and a gene encoding kanamycin resistance on a pET28a backbone. The plasmid was transfected into ClearColi by heat shock, then streaked for single colonies on kanamycin selective Luria Bertani (LB) agar plates. The identity of the plasmid was confirmed by whole plasmid sequencing (Plasmidsaurus). Colonies were picked after 2 days (ClearColi grows at half the rate of BL21 E.coli), and inoculated overnight at 37° C with shaking in a 10 mL LB-Kanamycin culture. Larger 250 mL cultures were inoculated and grown until they reached optical density 0.4. Cultures were induced with 250 mM Isopropyl ß-D-1-thiogalactopyranoside (IPTG) and grown overnight at 37° C after which time the bacteria were pelleted, and stored in a lysis buffer (50 mM NaH2PO4 at pH 7.4, and 300 mM NaCl) at -80°C until use. To purify protein, pellets were thawed, mixed thoroughly with a serological pipette, and then lysed in a French pressure cell press thrice. The lysate was clarified by centrifugation thrice, then incubated with Nickel NTA resin while shaking at room temperature for one hour. The resin was pelleted and washed thrice in wash buffer (20 mM sodium phosphate, 300 mM sodium chloride, 25 mM imidazole), and then bound enzyme was eluted thrice in the elution buffer (20 mM sodium phosphate, 300 mM sodium chloride, 250 mM imidazole). The eluants were pooled, buffer exchanged (to remove imidazole) into storage buffer (50 mM NaCl, 20 mM Tris-HCl, 1 mM EDTA, pH 7.5), assessed for protein concentration via BCA, run on an SDS page gel (**Fig S1**), then aliquoted into single use aliquots stored at -80° C until use.

#### Lectins and antibodies

Biotinylated Maackia Amurensis Lectin II (MAL II, Cat #:B-1265-1) and Cy5-conjugated Sambucus Nigra Lectin (SNA, Cat #: CL-1305-1) were purchased from Vector Labs. Human Contraception Antibody (HCA), was a gift from ZabBio. Recombinant human Galectin-1 was purchased from ACROBiosystems (Cat# 50-210-6461) and Pea savitum agglutinin (PSA-FITC) was purchased from Sigma (cat #L0770).

## Methods

### LC/MS-MS to confirm sequence of GvNanH2

In-gel digestion using trypsin/Lys-C and LC-MS/MS were used to confirm the protein sequence of the recombinant enzyme. LC-MS/MS analysis was carried out using a nanoAcquity UPLC (Waters Technology) interfaced to a Q-Exactive HF mass spectrometer (ThermoFisher Scientific.) Reversed-phase C-18 analytical and trapping columns were used with a 75 minute method with a gradient of 2% B to 40% B in 40 minutes, using 99% water/1% acetonitrile/0.1% formic acid as mobile phase A and 99% acetonitrile/1% water/0.1% formic acid as mobile phase B at a flow rate of 0.5 μl/min. Data-dependent tandem MS spectra were acquired in positive mode for the top 20 most abundant precursors. Full MS scans were acquired from *m/z* 350 to 2000 with a resolution of 60,000 using an automatic gain control target of 3e6 and maximum injection time (IT) of 100 ms. Dynamic exclusion (12 s) was enabled. Precursor ions were fragmented using a resolution of 15,000 with a maximum injection time of 50 ms and an automatic gain control value of 2e5 using higher energy collision-induced dissociation with a stepped normalized collision energy of 27 and 35 V. The MS/MS spectra were searched using Byonic against a fasta file containing the truncated NanH2 sequence and provided assignments for 81 peptides; these represented 83.2% of the predicted sequence (**Fig S2**).

### Sperm isolation

Fresh semen was acquired from healthy donors after obtaining written informed consent under the approved BUMC Institutional Review Board (IRB) protocol #H36843, and processed as described previously in Mausser et al. 2023. In brief, samples were processed within one hour of acquisition; motile sperm were separated by layering on a 90% ISolate density gradient (FUJIFILM Irvine Scientific; Santa Ana, CA, USA) and centrifugation at 300g for 20 minutes. The motile sperm pellet was resuspended in Multipurpose Handling Medium (MHM; FUJIFILM Irvine Scientific; Santa Ana, CA, USA). The concentration and motility of sperm in whole semen and isolated motile sperm were assessed using a Computer-Assisted Sperm Analysis system (CASA; Human Motility II software, CEROS II, Hamilton Thorne, Beverly, MA, USA). Samples with low concentrations (<10 million/mL) or motility (<40%) post-processing were not used for agglutination and complement experiments.

### Sperm sialidase treatment

Motile sperm were resuspended to a concentration of 30 million per mL in MHM. 40 μL of sperm suspension were incubated with 1 μL of enzyme at 37°C for the chosen time-points and enzyme concentration. All enzymes were thawed on ice, used fresh, and diluted into identical storage buffers (50 mM NaCl, 20 mM Tris-HCl, 1 mM EDTA, pH 7.5) to maintain identical media conditions and concentrations, even as enzyme concentration varied.

### Sialidase activity assay

Activity was confirmed by 4MU hydrolysis assay as previously described (Robinson et al. 2019). In brief, to measure sialidase activity and match levels of AUS, GvNanH2, PtNanH1 and PtNanH2 activity, a 4-methylumbelliferone-sialic acid assay was used. 20 μL of enzyme solution was diluted into 100 μL of 600 μM 4MU-sialic acid (Sigma Aldrich 69587), and immediately measured for fluorescence in a BioTek plate reader. The measurement was performed in a 96 well plate every 1 minute for 2 hours at 37°C (Ex/Em: 365nm/448nm, Gain=43). A standard curve of 2-fold dilutions in triplicate of 4MU starting at 150 μM and ending with 2.34 μM was used to convert arbitrary fluorescence units into μM 4MU hydrolysis per minute. The first 12 minutes of the reaction (<10% substrate consumption) was used as a linear portion of the curve to calculate activity. rNanH2 (produced recombinantly as described above) was used at a final concentration of at 0.27 ng/μL. AUS enzyme (NE BioLabs P0722L) at 20000 U/mL was diluted into 100 mM sodium acetate buffer (pH 5.5) for a final dilution of 1.8U/μL. Enzymes were thawed and kept on ice until use. *Prevotella* enzymes were tested for activity to match activity levels to those used for GvNanH2 in various assays (**Fig** S1).

### Normalization of sialidase activity between different enzymes and calculations of enzyme doses

To normalize activity of enzymes, matched 4-MU activity levels were used on the same number of sperm in our assays. In our studies, sperm were used at a concentration of 35 million per mL, with volumes of 40 uL per treatment condition. Our activity levels in the data are described as concentrations per total volume, as 0.88 uM 4-MU hydrolysis per minute per uL in the highest treatment condition. An additional way is to describe enzyme activity per million sperm, which is 25.1 4-MU hydrolysis per minute per million sperm.

### Inhibition of Sialidase Activity Assay

To measure activity of inhibited sialidase enzyme, activity assays were performed using purified NanH2 enzyme and N-Acetyl-2,3-dehydro-2-deoxyneuraminic acid inhibitor (DANA, Sigma Aldrich D9050). rNanH2 was used at a final concentration of 0.27 ng/μL. DANA was used at final concentrations ranging from 1 mM to 10 pM. Enzyme and inhibitor were added in triplicate in a black-bottom 96 well plate. The substrate, 4-methylumbelliferone-sialic acid (Sigma), was added immediately prior to reading the plate. Fluorescence activity was measured in a BioTek plate reader every 1 minute for 2 hours at 37°C (Ex/Em: 365nm/448nm, Gain=43). A standard curve of 2-fold dilutions in triplicate of 4MU starting at 150 μM and ending with 2.34 μM was used to convert arbitrary fluorescence units into μM 4MU.

### Endotoxin neutralization

To isolate the impact of endotoxin contamination in our experiments, enzymes were pretreated with 100 ug/mL of Polymyxin B (Sigma Aldrich, Cat #: 81271) for 10 minutes prior to testing in agglutination and complement assays as described. Polymyxin binds to the lipid A region on endotoxins and neutralizes their activity (Domingues et al 2012).

### Lectin flow cytometry

To measure the effect of sialidases on sperm surface glycopeptides, sperm were incubated with 160 U NanH2, 20 U AUS, or no treatment/heat-inactivated enzyme for 1 hour and were fixed in formaldehyde. After fixation, 5×10^4^ sperm were stained per condition at a final lectin concentration of 10 μg/mL for primary conjugates and 5 ug/mL for biotin conjugates for 30 minutes. Biotin conjugates were reacted with Neutravidin-Alexa Fluor 488 at 2 μg/mL for 45 minutes. Fluorescence on sperm was analyzed using a Cytek Aurora full spectral flow cytometer. Cells were gated by forward and side scatter and doublets were removed by gating the linear FSC-A vs FSC-H. Populations were compared by determining median fluorescence intensity for each condition in FlowJo software.

### Zeta potential measurements

Zeta potential was measured by diluting 20 μL of sperm at a concentration of 30 million per mL into 1 mL of distilled water immediately before measurement. A Malvern zetasizer instrument was used at the following settings: the diluent setting was 10mM NaCl, and seven measurements were taken per sample with no cooldown time between measurements.

### Sperm agglutination assay

Sperm were treated with sialidase as described above. Sperm with and without sialidase pretreatment were exposed to three sperm agglutinating agents: Human Contraception Antibody (HCA), and Galectin-1 and Pisum sativum agglutinin (PSA) lectins. A kinetic agglutination assay was performed as previously described (Mausser et al 2023.) to measure time to 95% agglutination.

### Complement-mediated sperm immobilization

Sperm were treated with sialidase as described, then exposed to human serum complement for 1 hour. The assay was performed as previously described (Mausser et al. 2023), except antibody treatment was excluded. In brief, motile sperm were quantified with CASA software after complement exposure. Heat-inactivated (Hi) sialidase enzyme and Hi complement were used as negative controls. A human complement inhibitor, compstatin (MedChemExpress, USA), was used as an additional negative control. Complement and compstatin were incubated together for 15 mins at room temperature and then sialidase-treated sperm were added as previously described. Classical complement-mediated sperm immobilization was measured with an identical protocol, but including human contraception antibody (HCA). As additional controls, sialidase activity was inhibited by addition of DANA (N-Acetyl-2,3-dehydro-2-deoxyneuraminic acid) at 1mM or Zanamivir at 1mM (MedChem Express, Cat. No.: HY-13210) to the maximum dose of enzyme (0.88 micromolar 4MU hydrolysis/min/uL).

### Human cervical mucus penetration assay

Cervical mucus samples were acquired after obtaining written informed consent with the BUMC Institutional Review Board (IRB) approved protocol #H41454. Midcycle ovulatory cervical mucus was collected from reproductive aged women using an endocervical pipelle (Aspirette Endocervical Pipelle, Cooper Surgical, Trumbull, CT, USA) within 48 hours of a positive ovulation test (Digital Ovulation Predictor Kit, Clearblue). Participants had regular menstrual cycles, were not taking any hormonal birth control, and refrained from intercourse for at least 48hrs prior to collection. The samples were stored at 4° C until use (within 5 days). Cervical mucus was diluted at a 1:3 ratio of mucus: PBS and aspirated into capillary tubes (Borosilicate Capillary Glass Slide, 0.30×3.0 mm, 50 mm, Electron Microscopy Sciences, Hatfield, PA, USA) using a squirrel feeder and syringe. One end of the tube was sealed using nail enamel, another end was filled with 10 μl of NanH2 sialidase or media alone as a control. The capillary tube was inserted horizontally in a 1.5ml microcentrifuge tube containing 80 μL of 20 x10^6^/mL motile sperm. The assay was performed as previously reported (Mausser et al, 2023). CASA images were taken at different points along the capillary tube to evaluate the number of motile sperm reaching distances of 1, 2, 3 and 4 cm.

### Sperm motility assay

Sperm motility was quantified over time using CASA software and specialized microscope slides with a shallow chamber depth. Sperm at 30 million per mL were treated with sialidase, incubated at 37C in a humidified chamber, and measurements were taken at specific time points in the CASA. Fields of view lacking spontaneous agglutinates were captured in this assay to enable accurate measurement of motility in the CASA software. At least 300 sperm were measured per condition to determine a percent motility corresponding to 3 to 7 fields of view per time point per condition. To measure hyperactivation parameters including curvilinear velocity, amplitude of lateral head displacement, and linearity for motile sperm, the average values calculated by the CASA software were plotted. In particular, these average values were calculated for motile sperm only, to exclude dead sperm with may skew the averages towards zero motility and hide the signal from hyperactivated motility.

### Sperm Live/Dead staining after complement treatment

Sperm were incubated with sialidase and serum complement, as described for the complement-mediated immobilization assay. Motility was measured with the Computer Assisted Sperm Analysis (CASA) software, then cells were stained with trypan blue (1:1) to confirm whether percent motility inversely correlated with the percentage of dead cells (trypan positive).

### Statistical analysis

Continuous variables with >2 conditions were analyzed by analysis of variance (ANOVA) or mixed-model analysis. A significant effect was followed by Holms-Sidak multiple-comparison tests, or a Tukey multiple-comparison test when Holms-Sidak could not be calculated. Variables were tested for normality by the Shapiro-Wilk test. If continuous variables were not normally distributed, they were log (natural)-transformed prior to analysis. For repeated measures ANOVA, the equality of variances of the pairwise differences between within-subject conditions was assessed through tests of sphericity (Greenhouse-Geisser epsilon and Huynh-Feldt epsilon). For 2-group/condition comparisons of continuous variables, unpaired or paired t-tests or Mann-Whitney U tests or Wilcoxon matched-pairs signed-rank tests comparing the slopes of linear regression lines were performed. Statistical significance was assumed when P<.05. Data analysis and graphing were performed with Prism, Version 9.4.1 (GraphPad Software, Inc, La Jolla, CA) software.

## Funding

This work was supported by the National Institutes of Health [P50 HD096957 to D.J.A and T32-A1007309 to S.D] and a Boston University Sexual Medicine Pilot Grant to J.M.

## Acknowledgements

We thank Warren and Amanda Lewis for generously providing the plasmids encoding the *Gardnerella* sialidases, and Emily Balskus, and her students Claire Xiang and Paula Pelayo for generously providing the *Prevotella* sialidase enzymes. We also thank Daniel Dempsey, Nader Rahimi, Bjoern Reinhard, and Joseph Zaia, and members of their lab(s), especially Laura Carretero Santos, Koustav Kundu and Margaret Downs, whose assistance and willingness to share equipment greatly aided this study. We thank Kevin Whaley for provision of HCA for these studies. We thank Shari Brezinsky from the BUMC Flow Cytometry Core Facility for outstanding technical help, and Jolinada Zhang and James Doud for contributing to motility and complement assays. Additionally, many thanks to Ashley Rebello, Seth Bloom, Cristina Zamora, Allysa Allen, and Matt Geib for providing advice regarding the study.

## Abbreviations

(BV): Bacterial vaginosis
(STI): sexually transmitted infection
(AUS): A. ureafaciens sialidase
(FRT): Female Reproductive Tract
(HCA): Human Contraception Antibody
(PSA): Pisum sativum agglutinin

## Data availability statement

The data that support the findings of this study are available from the corresponding author, [D.J.A], upon reasonable request.

## Author contributions

SD designed the research. SD, PS, and MF generated data. JP, SD and PS completed statistical analysis of the data. JP and JM obtained IRB approval and provided human samples for the study. CEC and DJA supervised the research and contributed to interpretation of the results. DJA and JM acquired funding for the study. SD took in the lead in writing the manuscript. All authors discussed the results and contributed to the final manuscript.

## Conflicting interests

The authors declare no conflicts of interest. The funders had no role in the design of the study; in the collection, analyses, or interpretation of data; in the writing of the manuscript; or in the decision to publish the results.

---

**Supplementary Figure 1:**
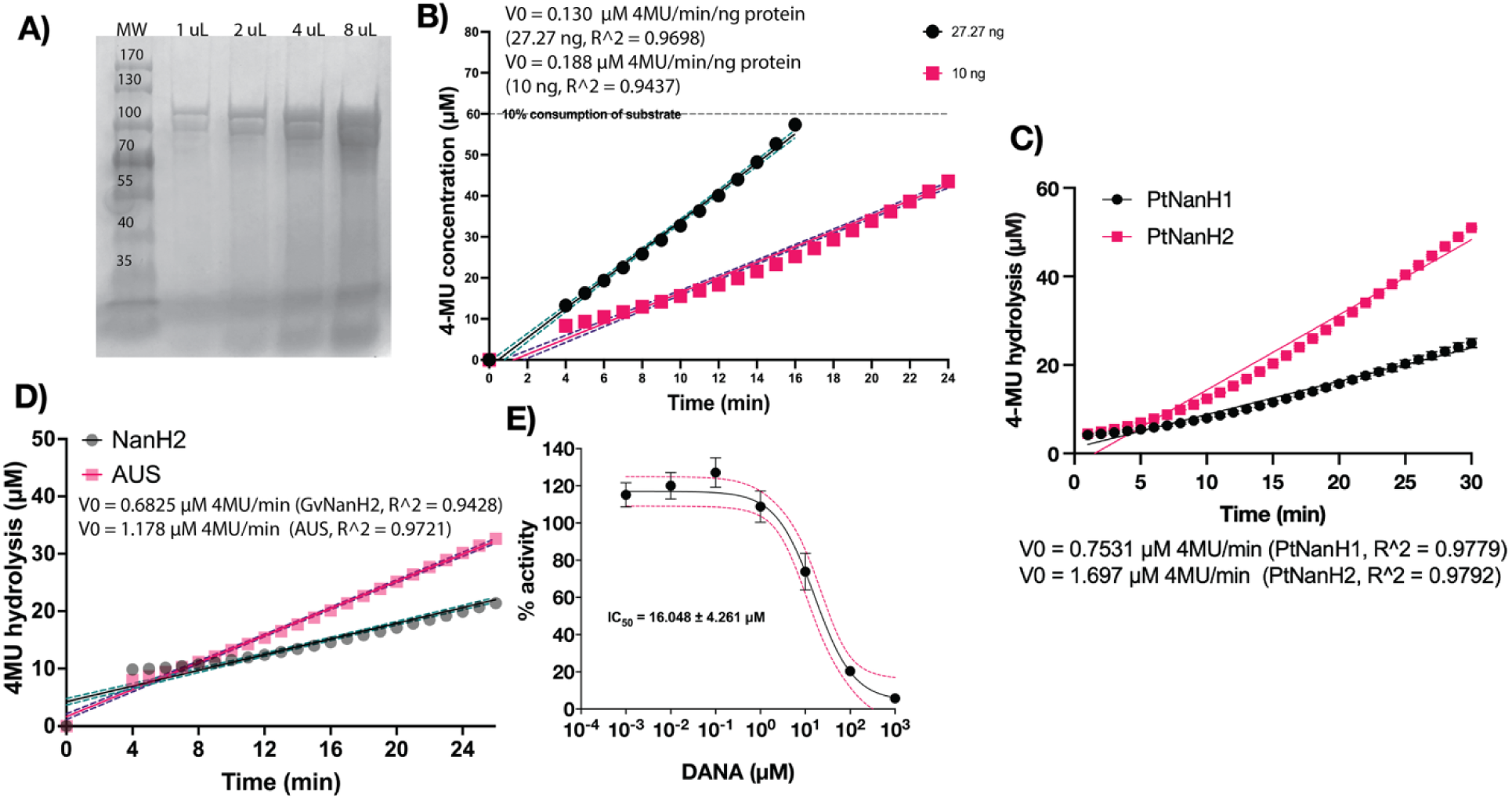
Production of GvNanH2 and characterization of Prevotella enzymes. A) SDS PAGE of purified GvNanH2. B) Activity determination of different concentrations of GvNanH2 with 4-MU hydrolysis assay. C) Activity determination of Prevotella enzymes stocks of with 4-MU hydrolysis assay. N= 3. D) Activity comparison to 5 units of commercially available AUS sialidase. N= 5 E) Inhibition of GvNanH2 by DANA, a sialidase inhibitor. 95% confidence intervals are represented by the pink dashed line. N = 3. Error bars represent standard error of the mean.

**Supplementary Figure 2:**
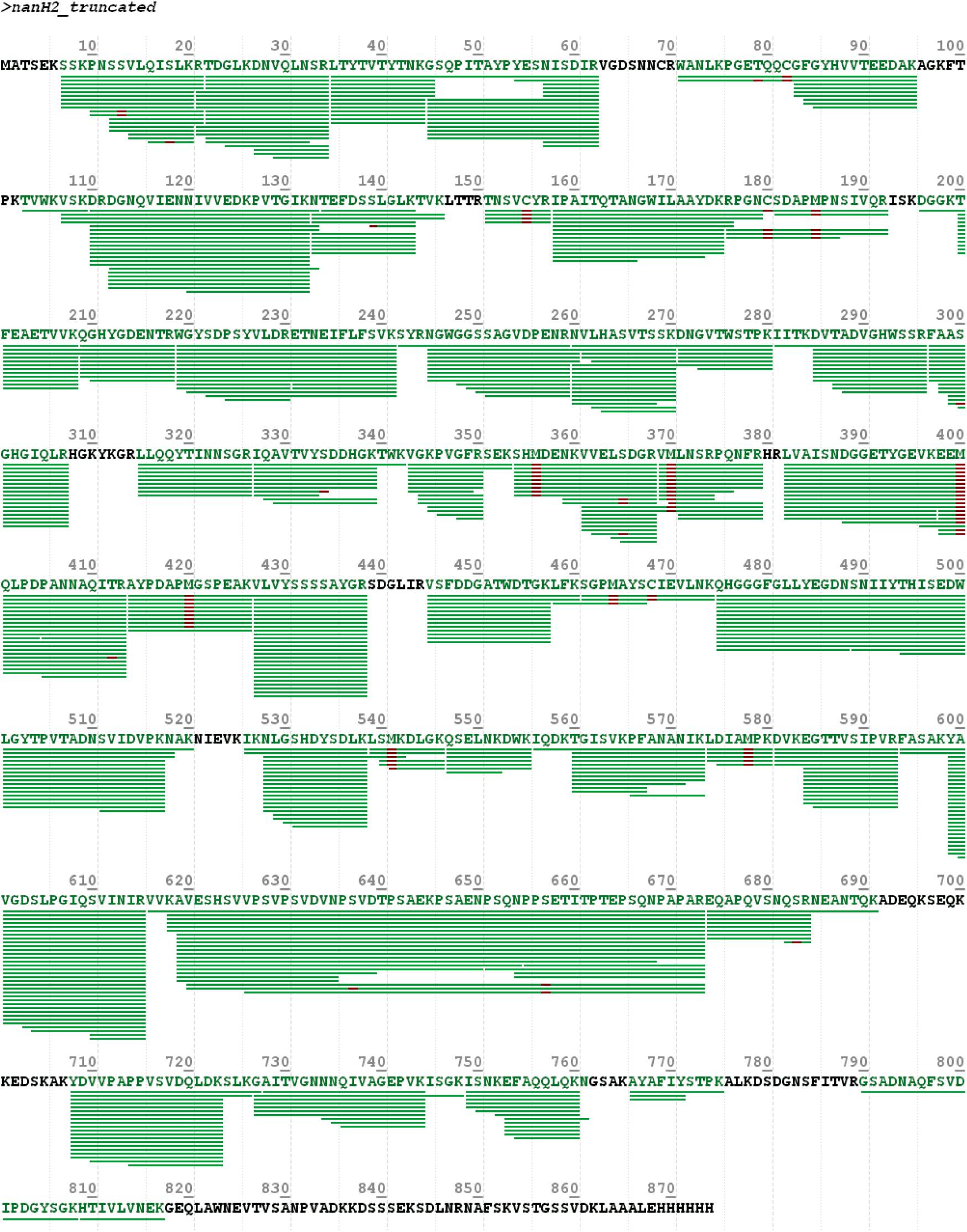
LC-MS/MS-based sequence coverage of in-gel digest confirms sequence of GvNanH2. The 75 Kda band from the SDS PAGE gel in Fig S1A was cut and digested with trypsin, and peptides were subjected to LC-MS/MS analysis to confirm the identity of the protein.

**Supplementary Figure 3:**
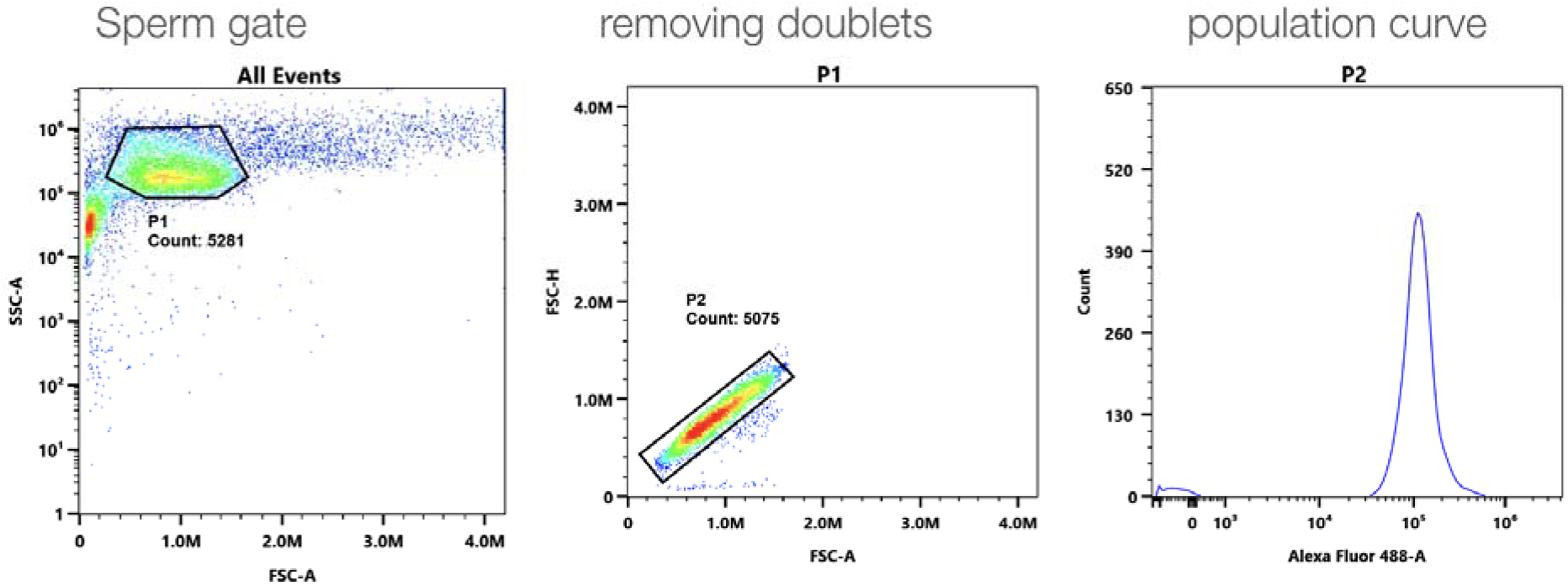
Flow cytometry gates for sperm. Sperm were fixed in formalin, stained with MAL-II, and analyzed by flow cytometry. Gates were selected to select sperm and remove doublets based on forward and side scatter.

**Supplementary Figure 4:**
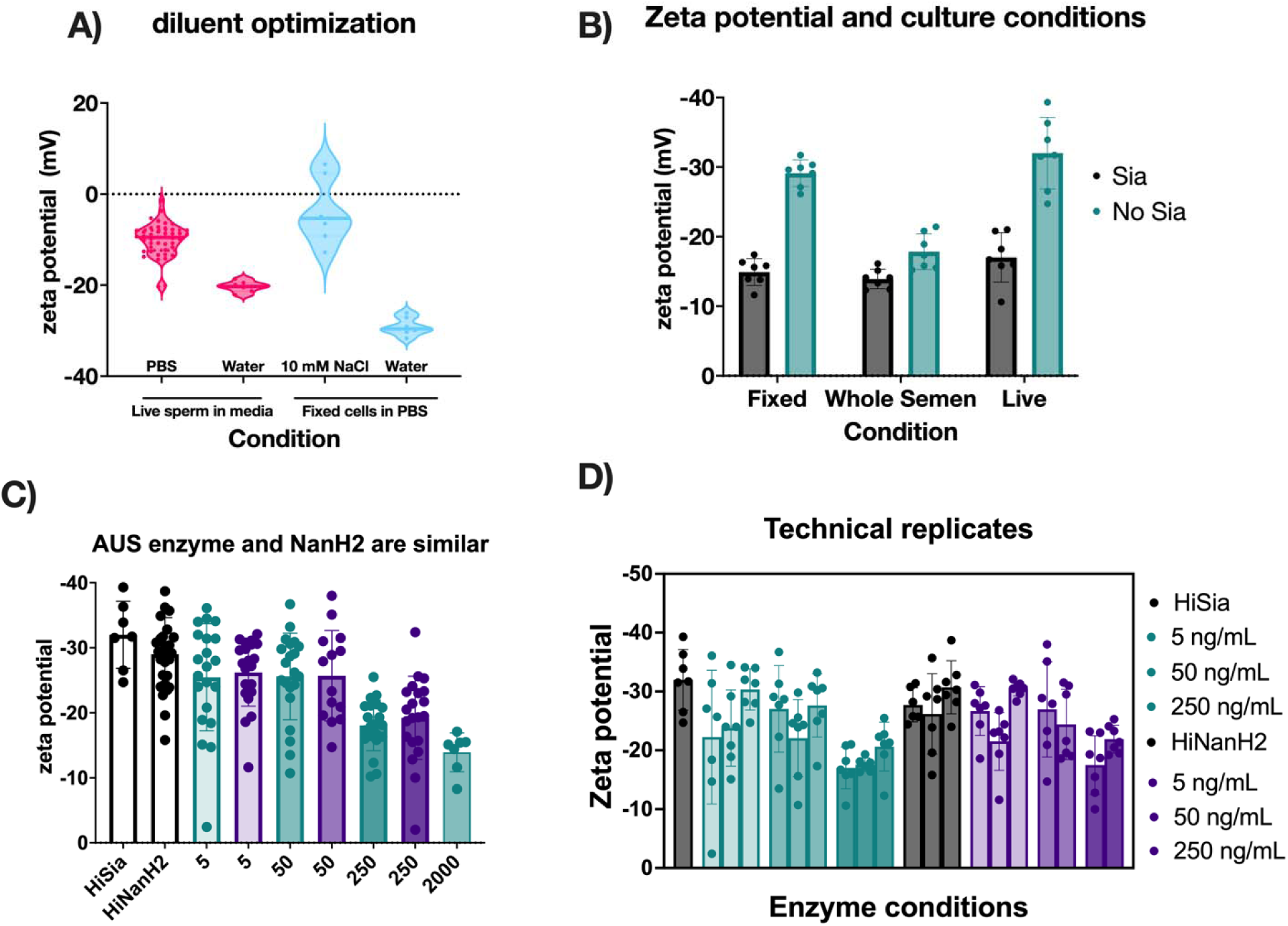
Zeta potential method development. A) Diluent optimization of reducing variability within technical replicates. B) Zeta potential and sperm culture conditions. C) Comparison of AUS (green) and NanH2 (purple). D) Biological replicates shown individually to show technical replicates within each sample, and highlight donor to donor variability. 2000ng/mL corresponds with 0.88 4-MU hydrolysis/min/uL.

**Supplementary Figure 5:**
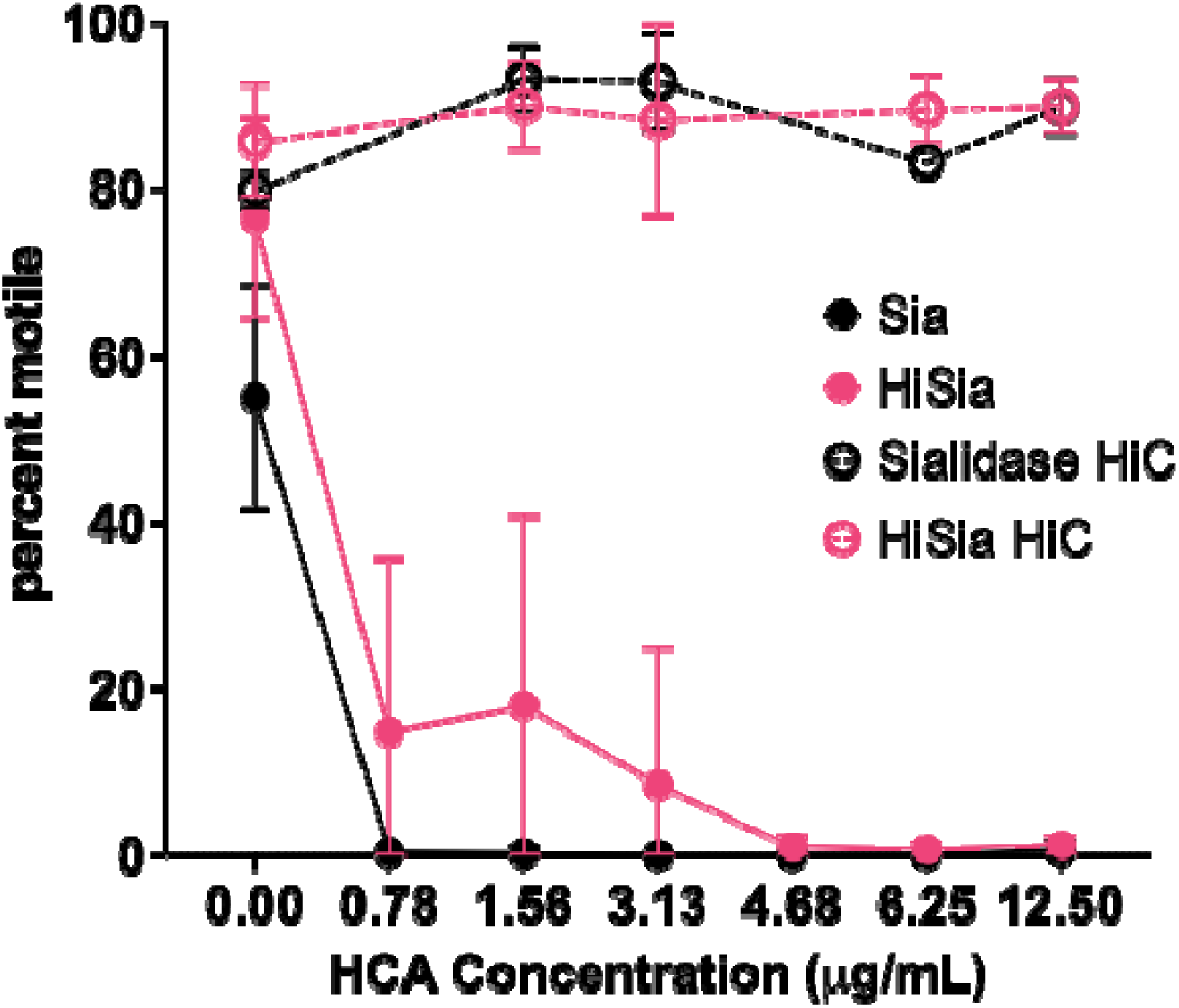
Human Contraception Antibody (HCA)-mediated complement immobilization of sperm is enhanced in the presence of sialidase. AUS sialidase at 20 units was incubated with sperm for 1 hour, then complement was added and incubated for 1 hour. Motility of these samples was quantified by CASA software. Each point represents the average of biological replicates and error bars represent standard deviations. A mixed effects analysis revealed that there was a statistically significant effect on motility due to the sialidase treatment and antibody dose (p<0.001), and post-hoc Holms-Sidak testing revealed a difference in the sialidase treatment conditions in the absence of antibody. N = 3 per biological replicate. Hi=heat inactivated.

**Supplementary Figure 6:**
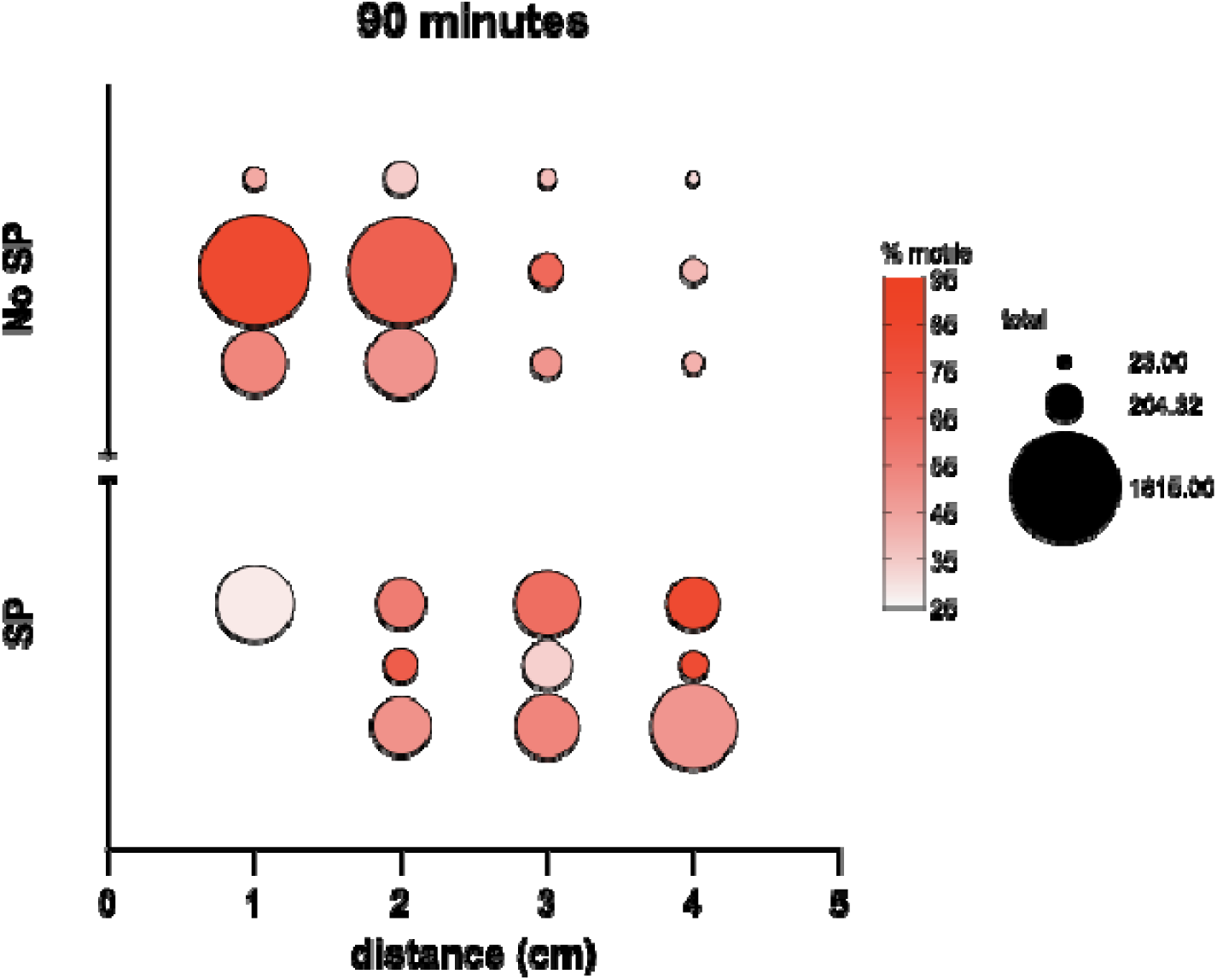
Effect of seminal plasma on cervical mucus transit of sperm. A cervical mucus transit assay was performed with one cervical mucus sample containing sialidase enzyme in the presence and absence of seminal plasma. The bubble size represents total sperm count at each point in the capillary tube, and the coloring represents the percent motility. Each point represents a technical replicate.

**Supplementary Figure 7:**
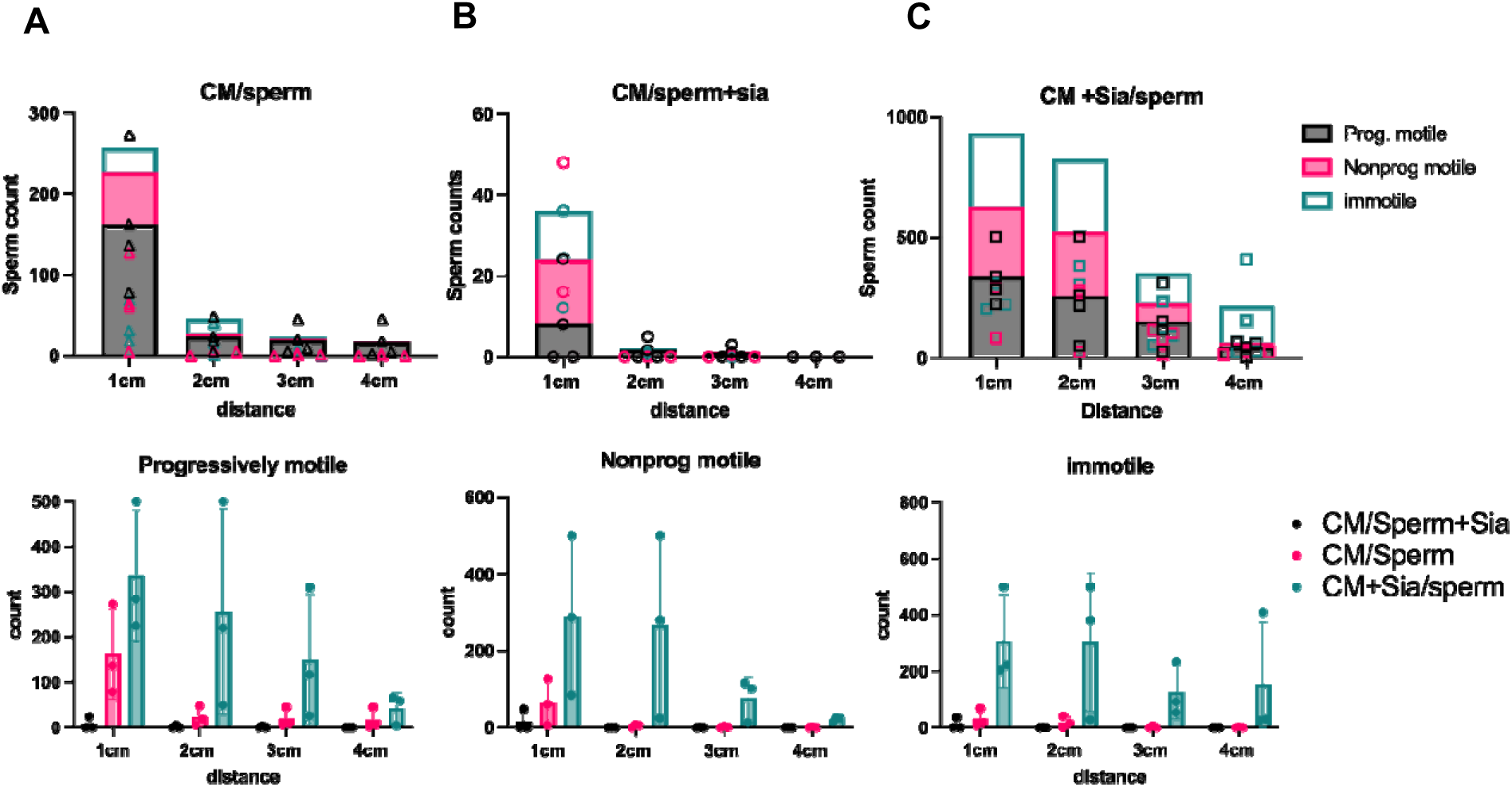
Method development for cervical mucus penetration. To optimize the cervical mucus transit assay, several variants of the assay were performed with a single cervical mucus sample. In A) no sialidase was added as a control condition. In B), sialidase was added to sperm samples only. In C), sialidase was per-mixed with cervical mucus. Each point represents a technical replicate. Data in the top row are stacked graphs representing both sperm counts and contribution of each motility category divided by experimental condition, and the bottom row is split by motility category to compare different experimental conditions.

**Supplementary Figure 8:**
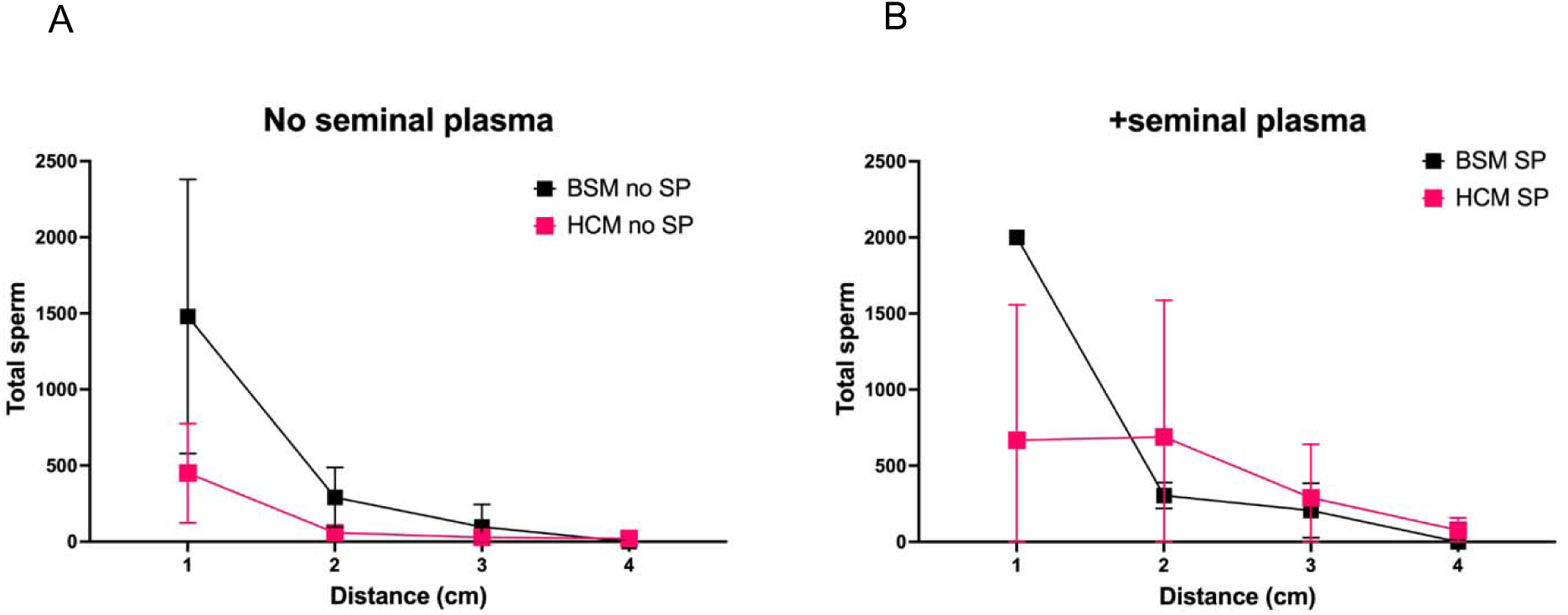
Comparison of human cervical mucin (HCM) and bovine submaxillary mucin (BSM) in sperm transit assay. Cervical mucus transit assay was performed with 1.5% bovine submaxillary mucin in PBS, to compare sperm transit with and without seminal plasma as part of assay development. Each point represents the average of three technical replicates, and error bars represent standard deviations. In the absence of seminal plasma, the overall sperm transit pattern was retained, but in the presence of seminal plasma, the pattern of sperm transit deviated between the bovine and human mucus.

**Supplementary figure 9:**
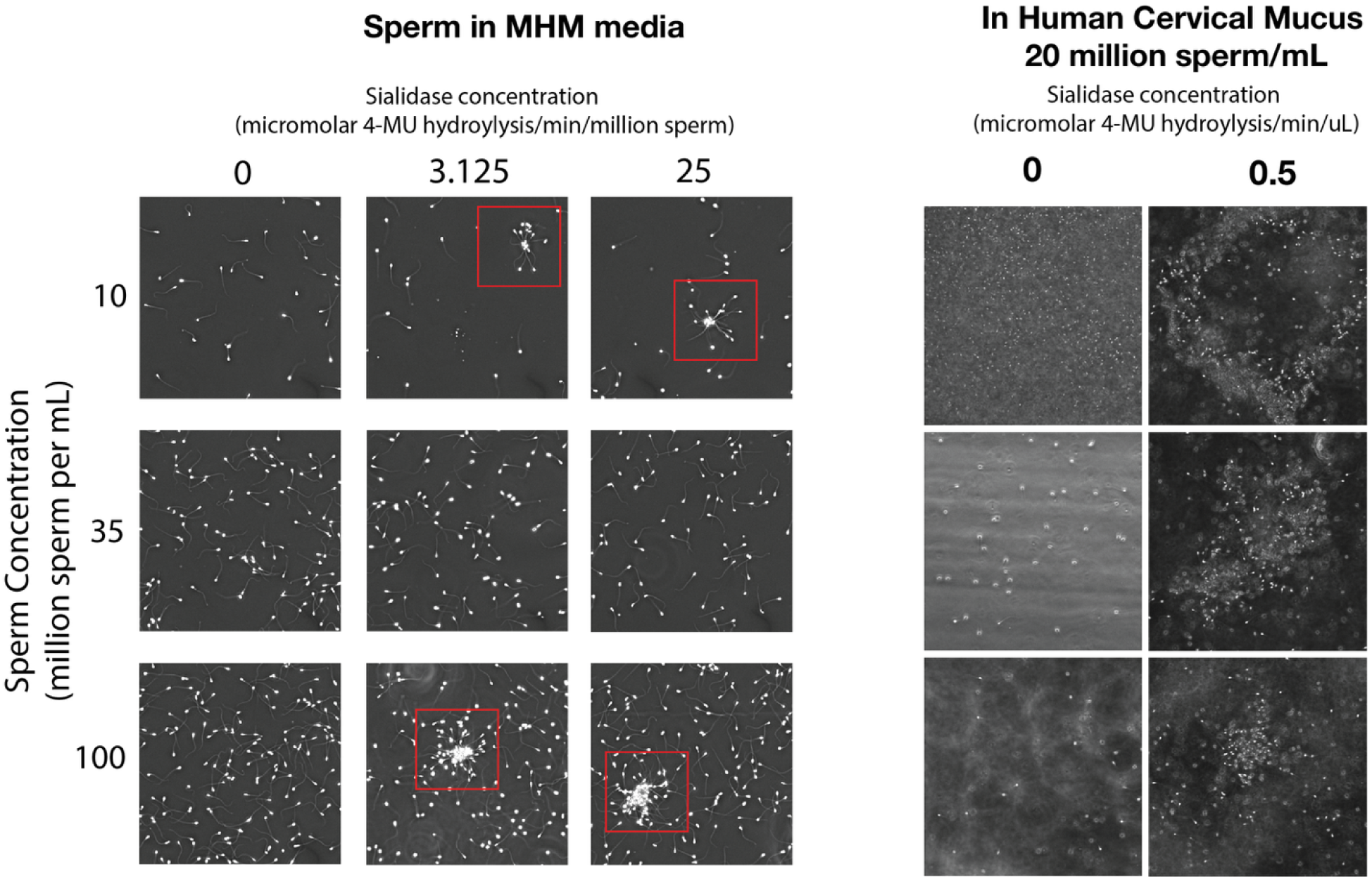
Spontaneous agglutination in the presence of sialidases. Motile sperm were exposed to PtNanH2 sialidase (0.88 4MU uM hydrolysis per min) for 1 hour and imaged for the presence of agglutinates. Some small clusters of sperm sticking together were observed. In cervical mucus, large agglutinates were sometimes observed in the presence of sialidase.

**Supplementary Figure 10:**
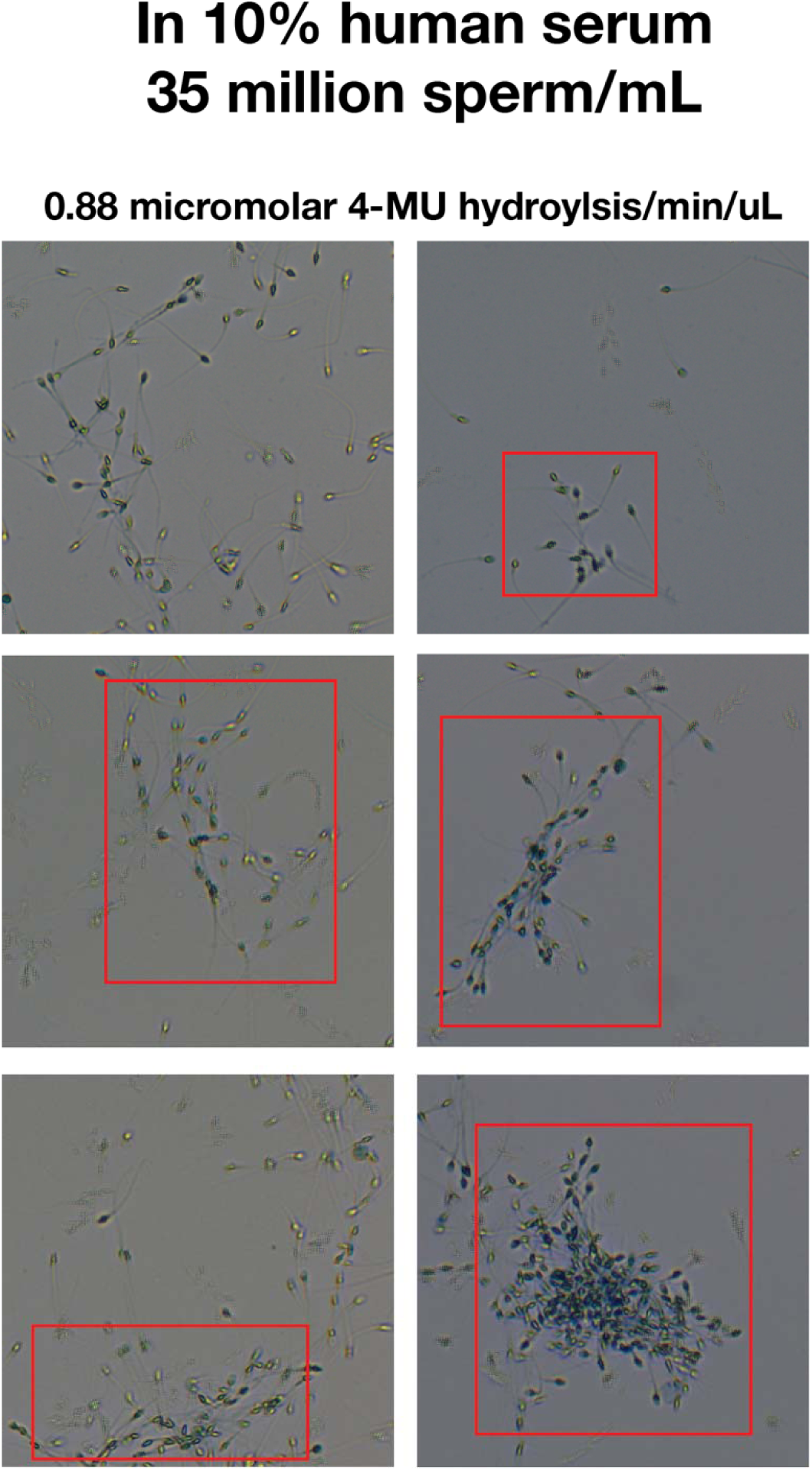
Representative images of sperm agglutination after treatment with NanH2 sialidase and complement for 1 hour. Sperm were stained with trypan blue to confirm cell death. Dead sperm were observed, along with motile live sperm (sometimes seen as tracks in the images due to movement).

## Notes

### Competing Interest Statement

The authors have declared no competing interest.

### Summary of Updates

Added additional data and revised introduction, results and discussion sections accordingly.

