## Supplementary Figures for "Sialidases derived from *Gardnerella vaginalis* and *Prevotella timonensis* remodel the sperm glycocalyx and impair sperm function"

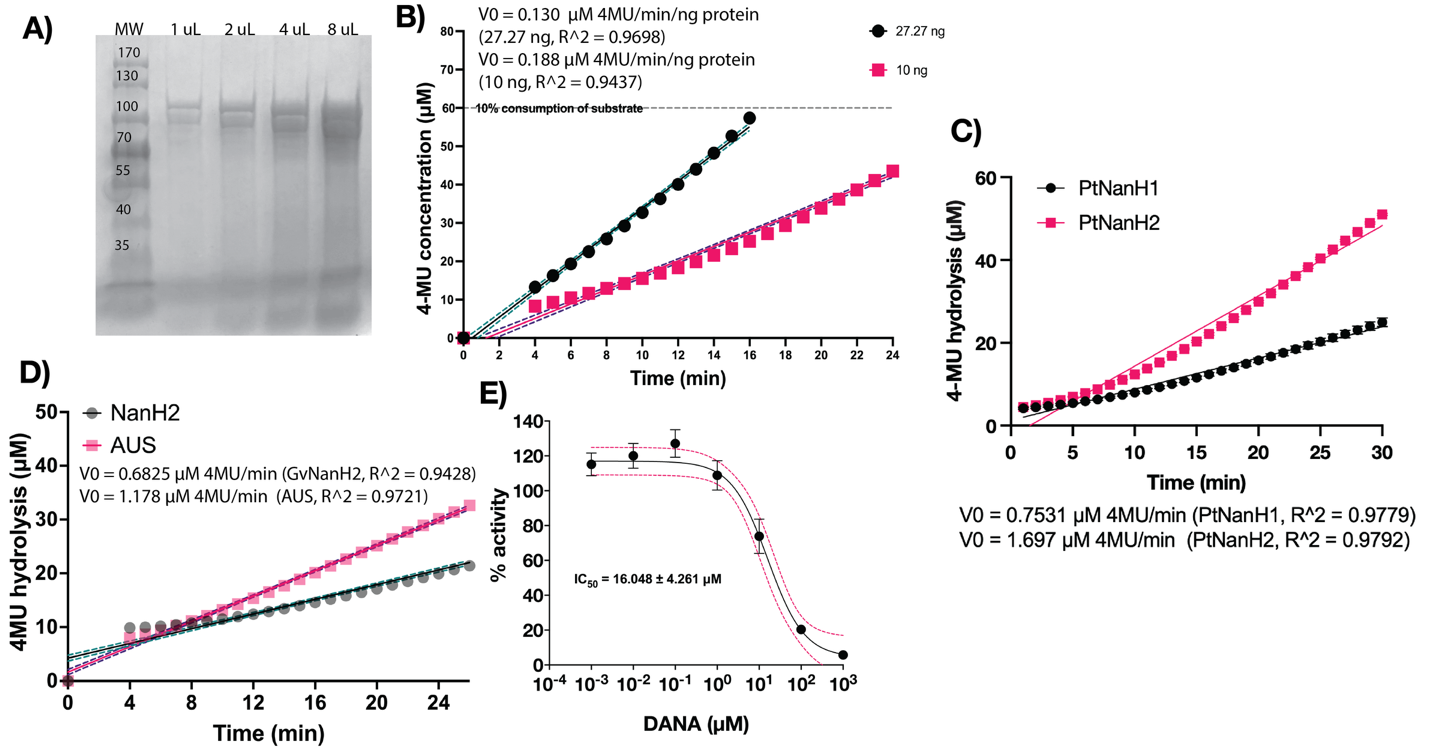

**Supplementary Figure 1: Production of GvNanH2 and characterization of Prevotella enzymes.** A) SDS PAGE of purified GvNanH2. B) Activity determination of different concentrations of GvNanH2 with 4-MU hydrolysis assay. C) Activity determination of Prevotella enzymes stocks of with 4-MU hydrolysis assay. N= 3. D) Activity comparison to 5 units of commercially available AUS sialidase. N= 5 E) Inhibition of GvNanH2 by DANA, a sialidase inhibitor. 95% confidence intervals are represented by the pink dashed line. N = 3. Error bars represent standard error of the mean.

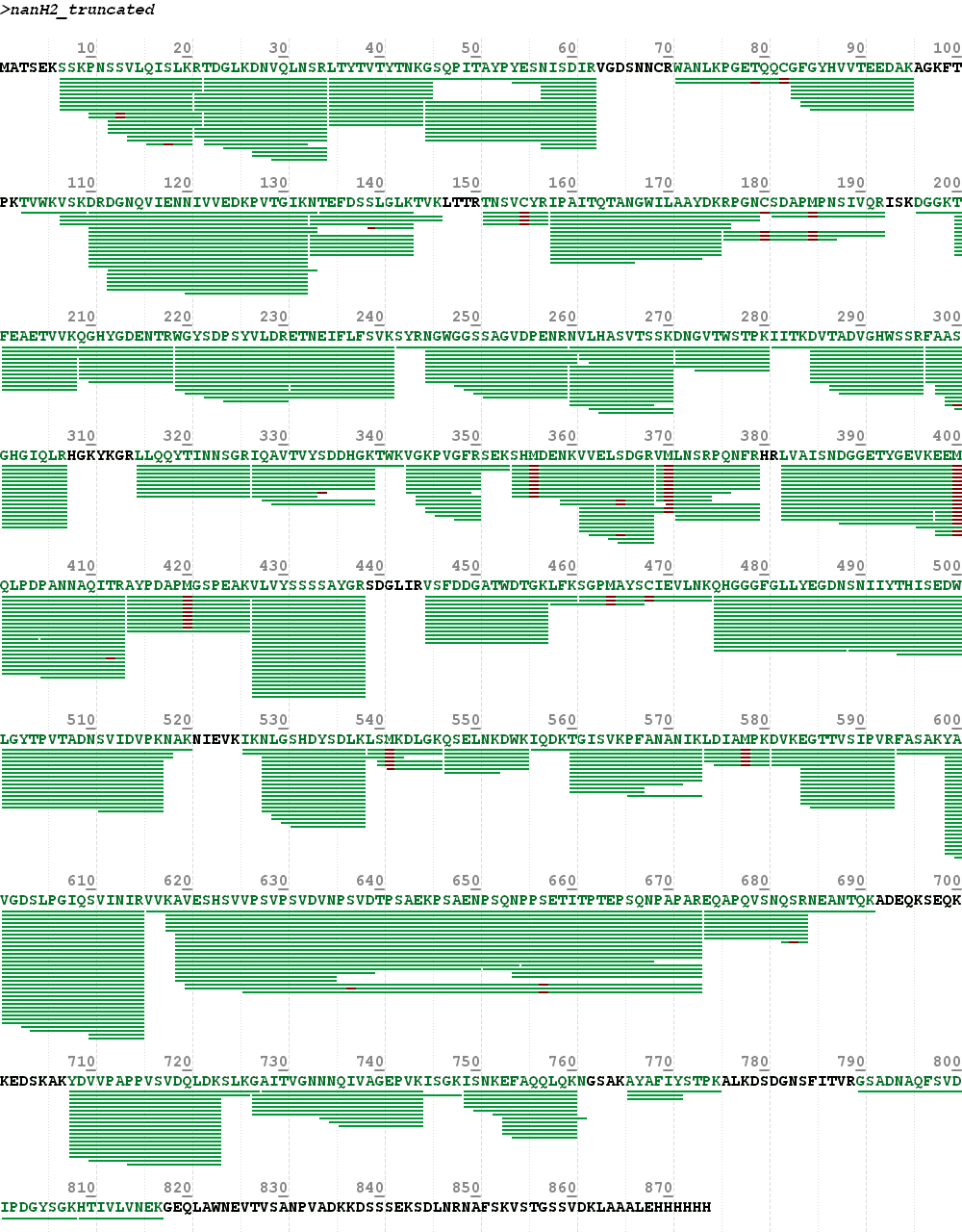

**Supplementary** **Figure** **2**: **LC-MS/MS- based sequence coverage of in-gel digest confirms sequence of GvNanH2.** The 75 Kda band from the SDS PAGE gel in Fig S1A was cut and digested with trypsin, and peptides were subjected to LC-MS/MS analysis to confirm the identity of the protein.

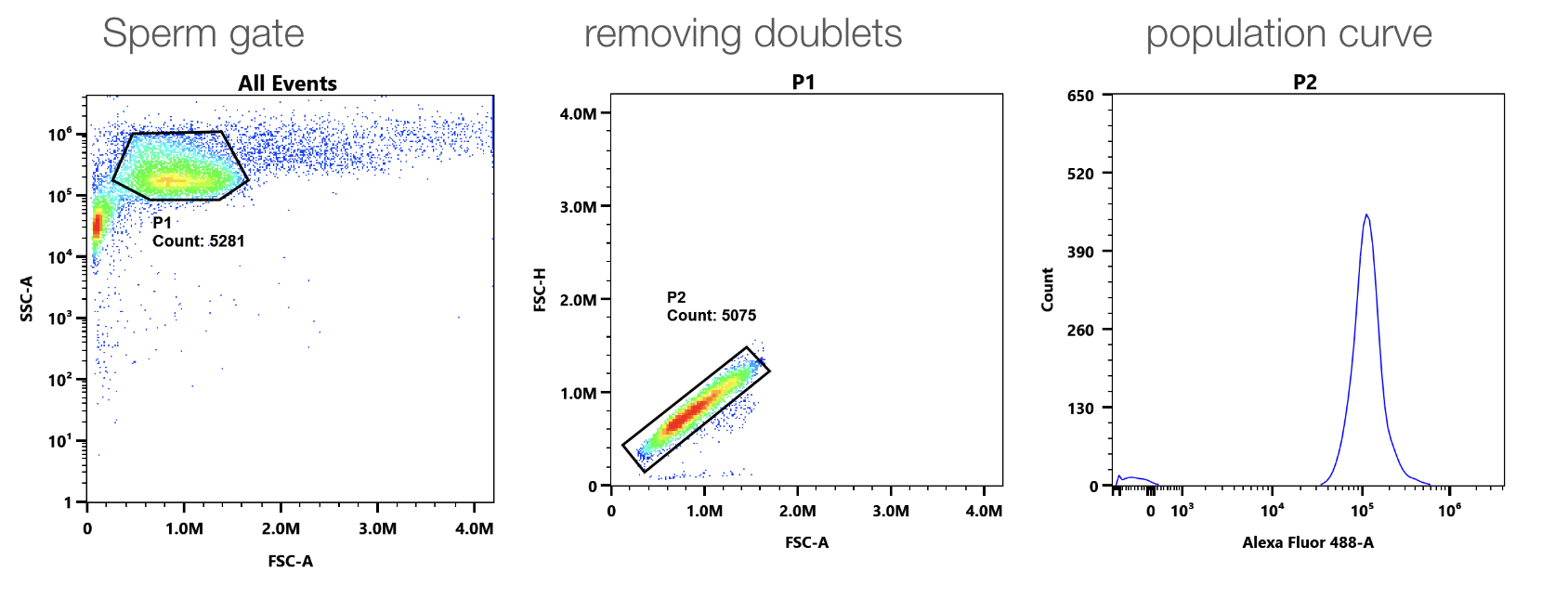

**Supplementary Figure 3: Flow cytometry gates for sperm.** Sperm were fixed in formalin, stained with MAL-II, and analyzed by flow cytometry. Gates were selected to select sperm and remove doublets based on forward and side scatter.

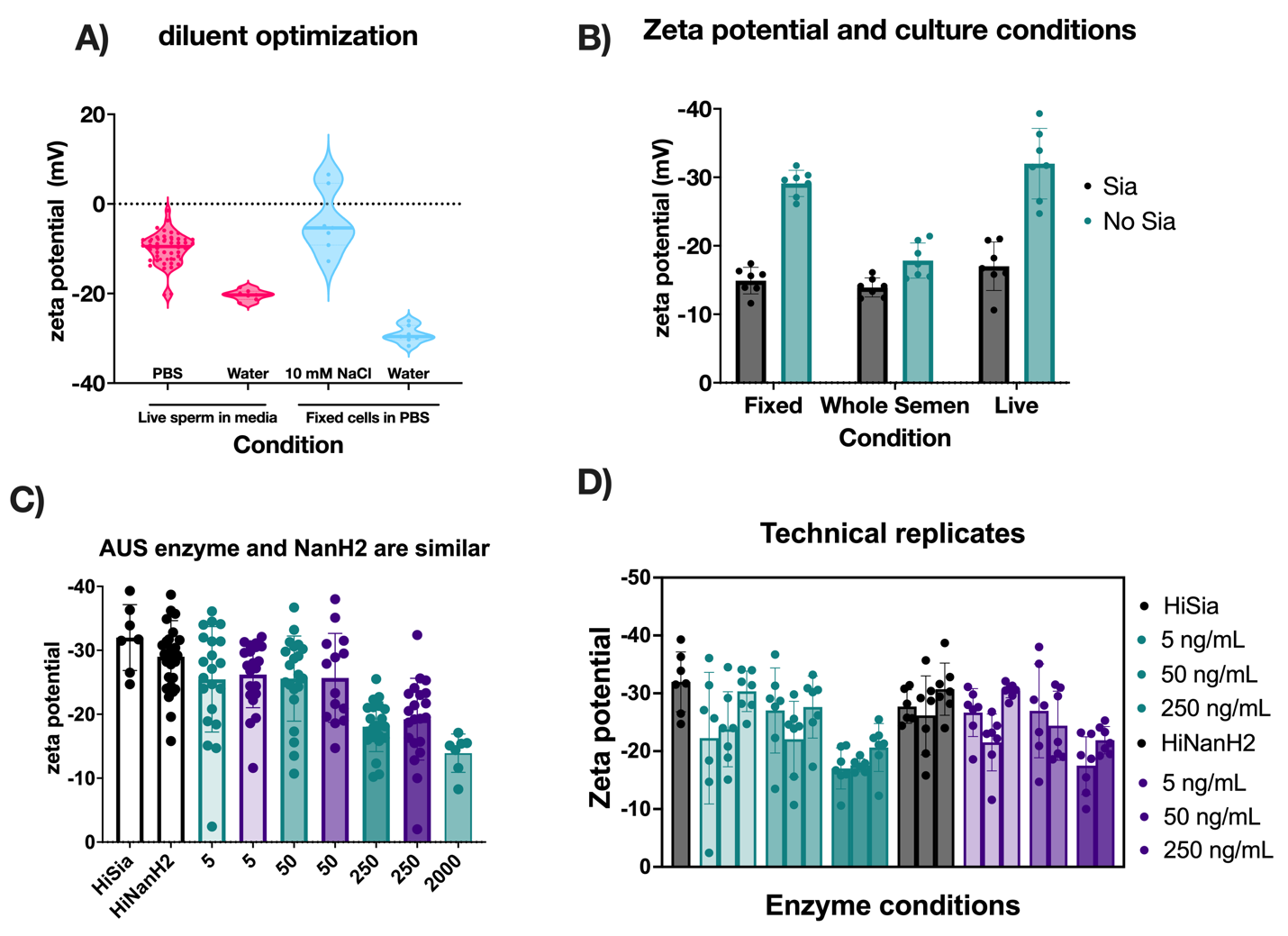

**Supplementary** **Figure** **4:** **Zeta** **potential** **method** **development**. A) Diluent optimization of reducing variability within technical replicates. B) Zeta potential and sperm culture conditions. C) Comparison of AUS (green) and NanH2 (purple). D) Biological replicates shown individually to show technical replicates within each sample, and highlight donor to donor variability. 2000ng/mL corresponds with 0.88 4-MU hydrolysis/min/uL.

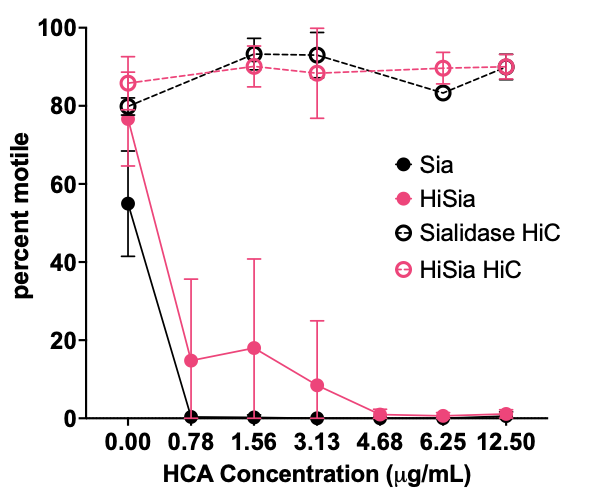

**Supplementary Figure 5: Human Contraception Antibody (HCA)-mediated complement immobilization of sperm is enhanced in the presence of sialidase.** AUS sialidase at 20 units was incubated with sperm for 1 hour, then complement was added and incubated for 1 hour. Motility of these samples was quantified by CASA software. Each point represents the average of biological replicates and error bars represent standard deviations. A mixed effects analysis revealed that there was a statistically significant effect on motility due to the sialidase treatment and antibody dose (p<0.001), and post-hoc Holms-Sidak testing revealed a difference in the sialidase treatment conditions in the absence of antibody. N = 3 per biological replicate. Hi=heat inactivated.

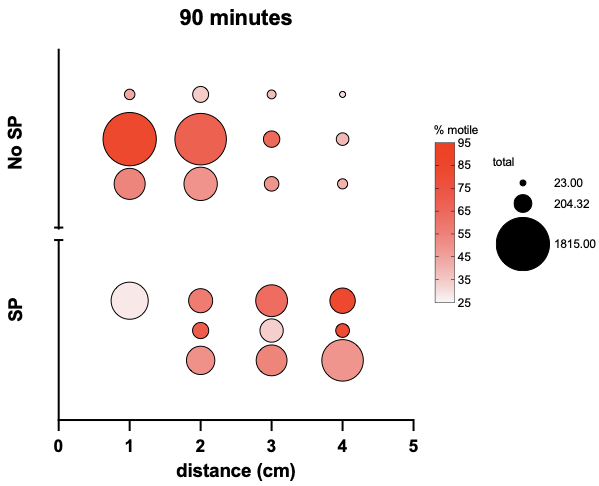

**Supplementary Figure 6: Effect of seminal plasma on cervical mucus transit of sperm.** A cervical mucus transit assay was performed with one cervical mucus sample containing sialidase enzyme in the presence and absence of seminal plasma. The bubble size represents total sperm count at each point in the capillary tube, and the coloring represents the percent motility. Each point represents a technical replicate.

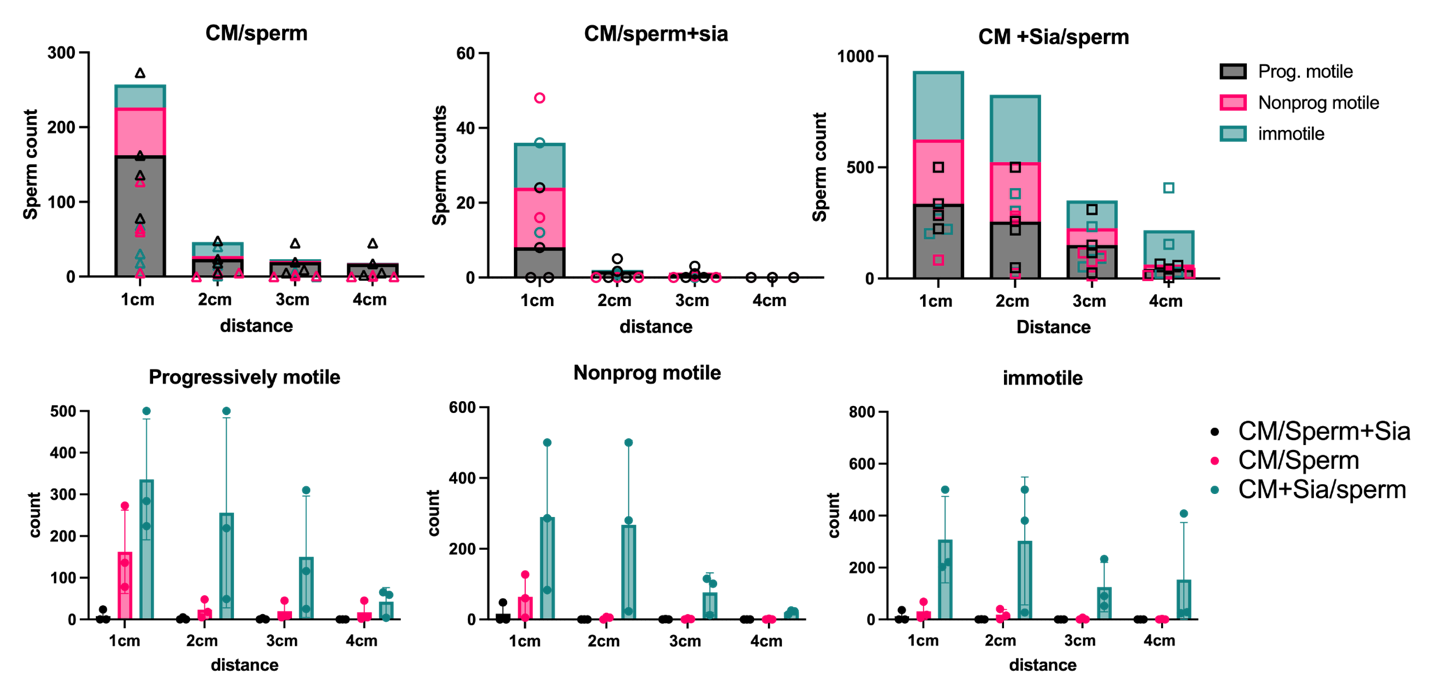
**Supplementary Figure 7: Method development for cervical mucus penetration** To optimize the cervical mucus transit assay, several variants of the assay were performed with a single cervical mucus sample. In A) no sialidase was added as a control condition. In B), sialidase was added to sperm samples only. In C), sialidase was per-mixed with cervical mucus. Each point represents a technical replicate. Data in the top row are stacked graphs representing both sperm counts and contribution of each motility category divided by experimental condition, and the bottom row is split by motility category to compare different experimental conditions.

**C**

**B**

**A**

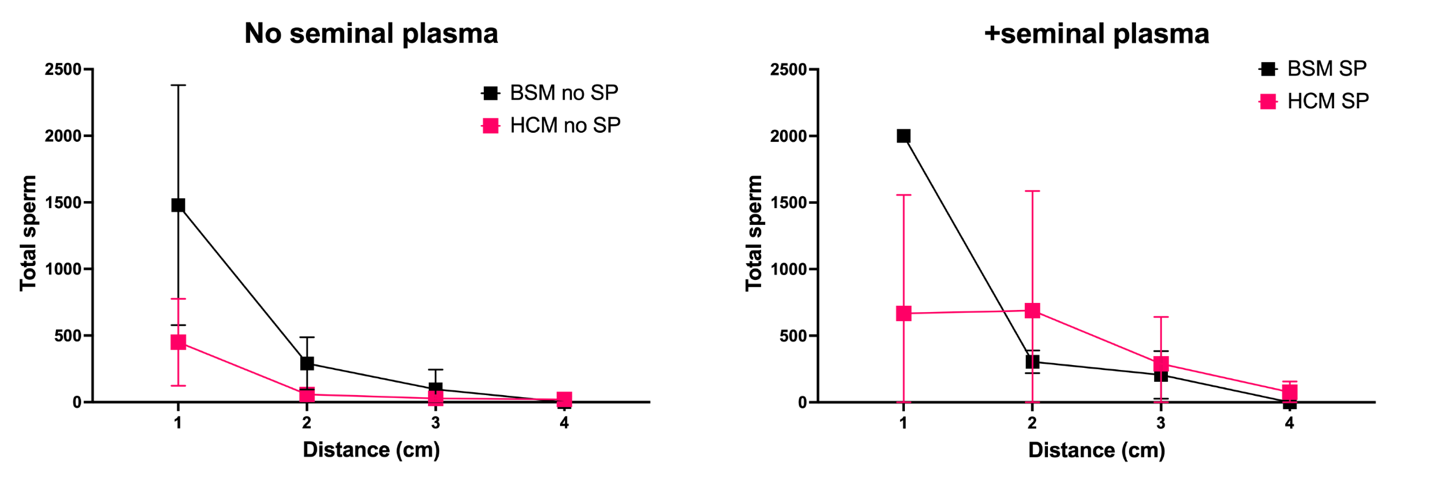

B

A

**Supplementary Figure 8: Comparison of human cervical mucin (HCM) and bovine submaxillary mucin (BSM) in sperm transit assay.** Cervical mucus transit assay was performed with 1.5% bovine submaxillary mucin in PBS, to compare sperm transit with and without seminal plasma as part of assay development. Each point represents the average of three technical replicates, and error bars represent standard deviations. In the absence of seminal plasma, the overall sperm transit pattern was retained, but in the presence of seminal plasma, the pattern of sperm transit deviated between the bovine and human mucus.

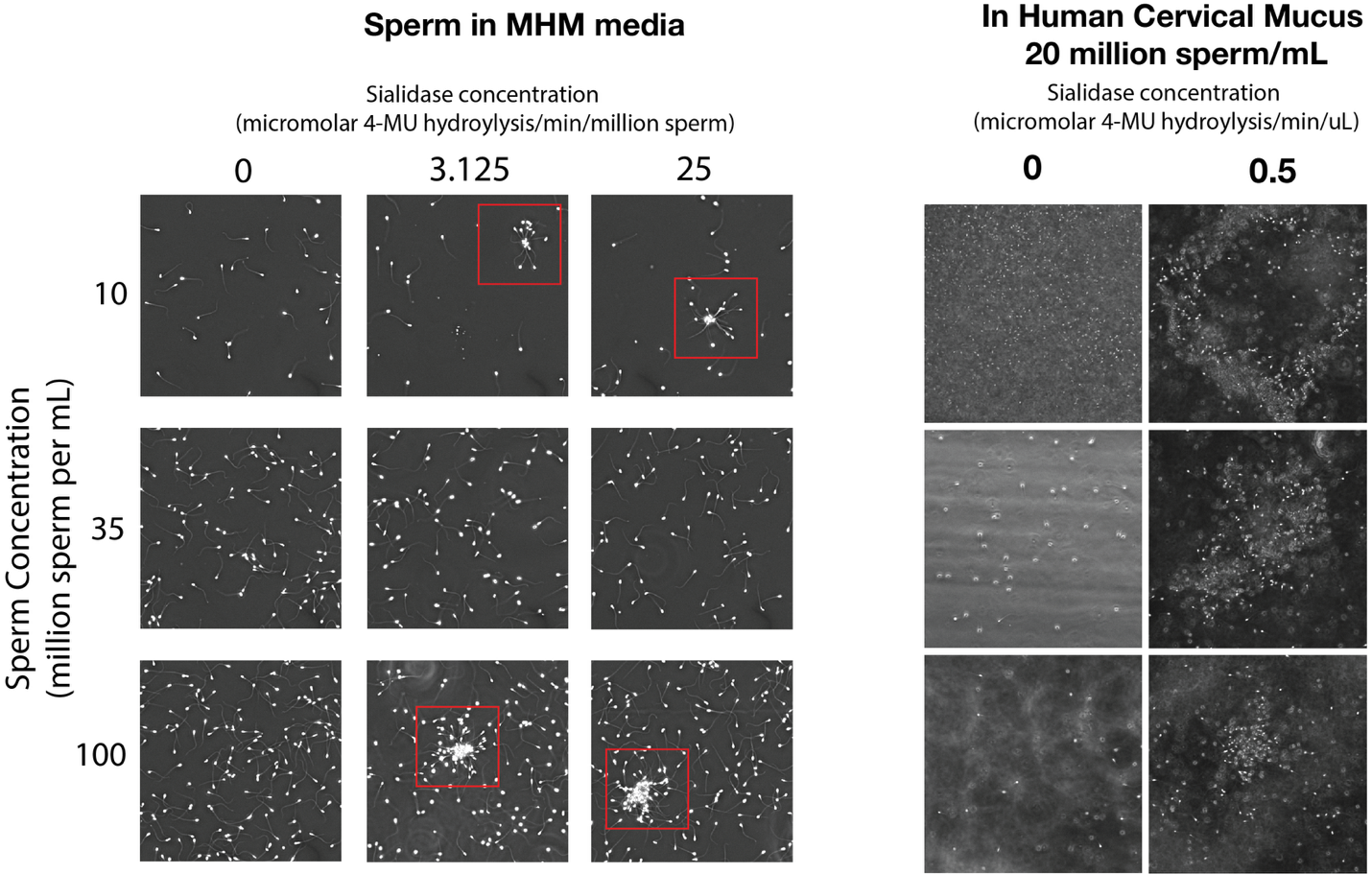

**Supplementary figure 9:** Spontaneous agglutination in the presence of sialidases. Motile sperm were exposed to PtNanH2 sialidase (0.88 4MU uM hydrolysis per min) for 1 hour and imaged for the presence of agglutinates. Some small clusters of sperm sticking together were observed. In cervical mucus, large agglutinates were sometimes observed in the presence of sialidase.

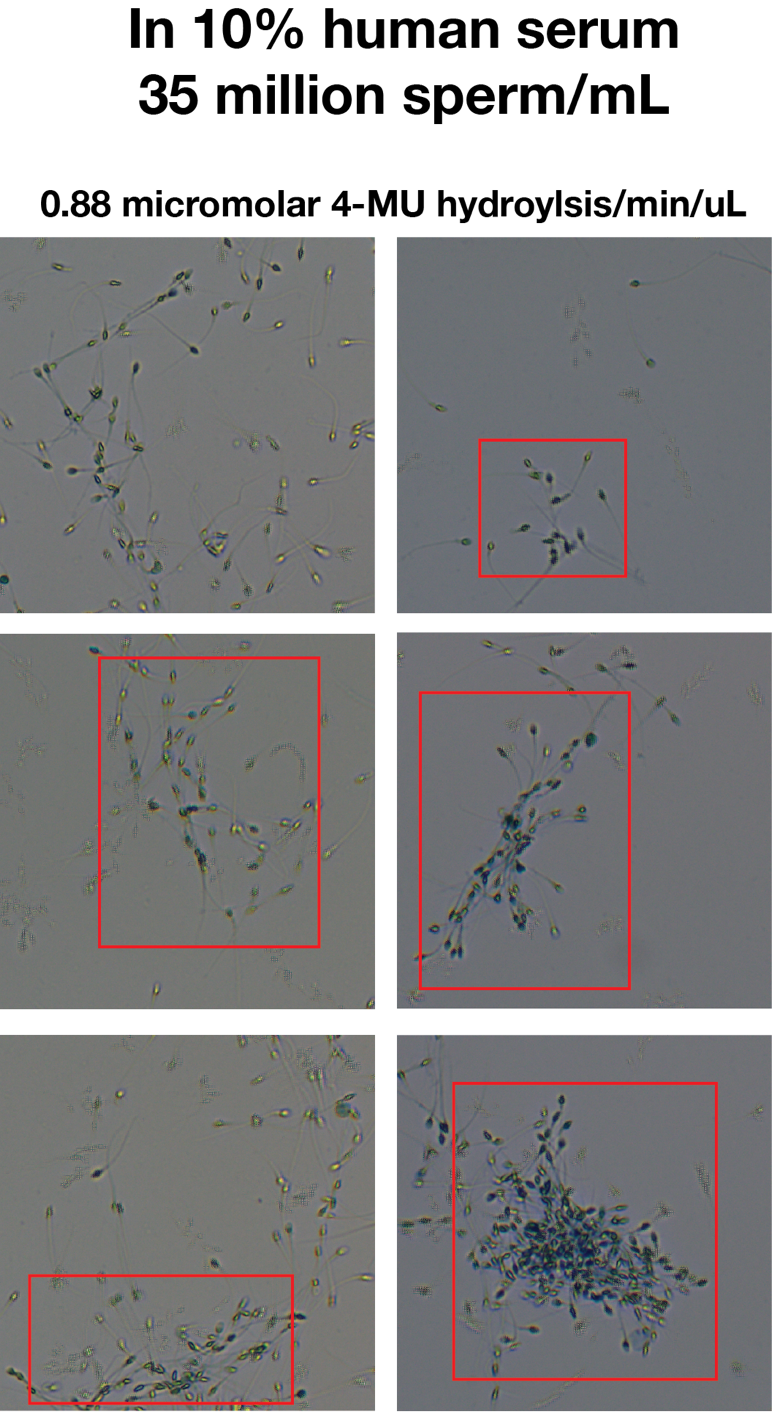

**Supplementary Figure 10:** Representative images of sperm agglutination after treatment with NanH2 sialidase and complement for 1 hour. Sperm were stained with trypan blue to confirm cell death. Dead sperm were observed, along with motile live sperm (sometimes seen as tracks in the images due to movement).
